# The Calcium Pump ATP2B1/PMCA1 Regulates CNS Vascular Development by Facilitating Norrin- and WNT7A/B-induced Frizzled4 Signaling

**DOI:** 10.1101/2025.07.17.664964

**Authors:** Ha-Neul Jo, Elizabeth Kiffmeyer, Chi Zhang, Lingling Zhang, Emmanuel Odame, Quynh Chau Dinh, Jacklyn Levey, Miranda Howe, Kyle J. Roux, Klaus-Dieter Fischer, Zhe Chen, Harald J. Junge

## Abstract

Frizzled4 (FZD4) is a receptor for Norrin and WNT7A/B ligands, is expressed in endothelial cells (ECs), is required for endothelial blood-central nervous system (CNS) barrier function as well as CNS angiogenesis, and transduces β-catenin-dependent signaling. Despite its fundamental importance in neurovascular biology, including as a drug target, the molecular mechanisms governing FZD4 regulation remain poorly understood. Here, we employed proximity biotinylation to identify proteins that regulate FZD4. We identified ATPase Plasma Membrane Ca²⁺ Transporting 1 (ATP2B1, also known as PMCA1) as a FZD4 proximity interactor. Functional analyses revealed that ATP2B1 depletion increased EC Ca^2+^, activated NFAT, and significantly attenuated Norrin/Frizzled4 signaling. Endothelial-specific *Atp2b1* deletion caused CNS vascular phenotypes consistent with compromised Norrin/Frizzled4 signaling. This study identifies ATP2B1 as a novel regulator of Norrin- and WNT7A/B-induced FZD4 signaling and suggests that in pathological contexts with elevated EC Ca^2+^-levels, EC function may be modulated by suppression of β-catenin-dependent signaling.

## INTRODUCTION

CNS ECs participate in angiogenesis under the control of CNS-specific mechanisms and are essential for the function of the blood–brain barrier and inner blood–retina barrier (BBB and BRB). In the endothelial barriers, ECs contribute to selective transport, regulation of the neural milieu, neuroimmune modulation as well as protection from toxins and pathogens. Defects in the BBB and BRB contribute to the onset or progression of diseases such as stroke, neurodegenerative disorders, retinopathies, and macular edema ^1, 2, 3, 4, 5^. A key pathway for inducing and maintaining the BBB and BRB and for enabling CNS angiogenesis is β-catenin-dependent signaling transduced by Frizzled and LRP5/6 receptors in ECs ^6^. β-catenin-dependent signaling in ECs can be induced by Norrin (gene symbol NDP) or WNT7A/B. Norrin has major functions in the retina and cerebellum ^7, 8, 9^ and WNT7A and B carry out major functions in the developing forebrain and spinal cord ^10, 11^. In many mature brain regions, Norrin and WNT7A/B function in parallel ^12^. Among the ten Frizzled (FZD1-10) receptors, FZD4 serves as the sole Frizzled receptor for the Norrin protein ^13^ and as one of the receptors for WNT7A/B ^14^. FZD4 associates with additional receptor complex components: Tetraspanin12 (TSPAN12) is a Norrin co-receptor that amplifies signaling in a ligand specific manner ^15, 16, 17^. The receptor complex components G protein-coupled receptor 124 (GPR124) and Reversion-inducing cysteine-rich protein with Kazal motifs (RECK) amplify WNT7A/B signals ^14, 18, 19, 20, 21, 22, 23, 24, 25, 26, 27, 28^. The causal relationship between impaired Norrin/Frizzled4 signaling and retinal hypovascularization in Norrie disease, osteoporosis-pseudoglioma syndrome, and familial exudative vitreoretinopathy (FEVR) underscores the critical role of this pathway in retinal vascular development ^29^. Retinal Norrin/Frizzled4 signaling is regulated by glutamatergic neuronal activity ^30^. In the brain, loss of β-catenin-dependent signaling in ECs is implicated in several neurological conditions including ischemic stroke ^31^. Loss of Norrin/Frizzled4 signaling in cerebellar ECs creates a permissive environment for medulloblastoma ^32^. Together, these studies establish critical functions of FZD4 in CNS vascular biology.

Given the pivotal role of FZD4 in CNS endothelium, this receptor has emerged as a target for experimental therapies currently in clinical development ^33^. The pharmacodynamic effects of Norrin-mimetics have been characterized in preclinical studies ^34, 35, 36, 37, 38, 39, 40^. Although FZD4 has garnered significant attention in neurovascular biology, the mechanisms governing its regulation in ECs are not well understood.

In this study, we identify ATPase plasma membrane Ca²⁺ transporting 1 (ATP2B1 aka PMCA1) as a novel regulator of Norrin- and WNT7A/B-induced FZD4 signaling in CNS ECs. As a plasma membrane calcium extruder pump, ATP2B1 maintains calcium homeostasis within cells and influences the shape and duration of Ca^2+^-transients ^41^.

The importance of these roles is underscored by the lethality resulting from its systemic deletion in mice ^42^. In addition to ATP2B1, the molecular repertoire for Ca^2+^-clearance in various cell types includes the paralogs ATP2B2-4, the Na^+^/Ca^2+^ exchanger family (NCX) and the Sarcoplasmic/Endoplasmic Reticulum Calcium ATPase (SERCA) family, which transports Ca^2+^ into the ER. It is thought that cytosolic nanodomains with low Ca^2+^-concentration in immediate proximity to ATP2B1 modulate local Ca^2+^-dependent processes, possibly supported by physical confinement, e.g., a plasma membrane invagination. The association of ATB2B1 with scaffold proteins may help to recruit receptors into low Ca^2+^-nanodomains in the vicinity of ATP2B1 ^43^. ATP2B1-4 have been studied in various excitable cell types, including sensory neurons, smooth muscle cells, and cardiomyocytes ^44^ as well as various non-excitable cells such as lymphocytes ^45, 46,47^. Ca^2+^ in ECs is involved in responses to signals that modulate vasodilation, angiogenesis, vascular permeability, thrombogenesis, diapedesis or vascular inflammatory responses, including ATP, VEGF-A, bradykinin, histamine, thrombin and shear stress ^48, 49, 50, 51, 52, 53^. Few studies have investigated the roles of ATP2B1–4 in ECs. ATP2B1 has been studied in relation to HUVEC migration ^54^ and nitric oxide production ^55^, while ATP2B4 has been examined for its role in VEGF-A–induced angiogenesis ^56^. Here, we identify ATP2B1 as a proximity interactor of FZD4 and elucidate a novel role in Norrin- and WNT7A/B-induced FZD4 signaling using cell-based and mouse genetic approaches.

## RESULTS

### Proximity interaction screens with a FZD4 bait

To identify novel regulatory mechanisms in FZD4 signaling, we performed proximity labeling. This approach enables detection of proximity interactions within approximately 10 nm radius in living cells using BioID, a promiscuous form of the bacterial BirA biotin ligase ^57^. We used V5-FZD4-BioID as bait and GFP-BioID as a control. V5-FZD4-BioID may identify proximity interaction during intracellular trafficking through the secretory pathway, at the plasma membrane, or during endo-lysosomal trafficking (Fig. 1A). The functionality of the V5-FZD4-BioID fusion protein was tested by examining V5-FZD4-BioID subcellular localization in HeLa cells. In live HeLa cells subjected to flag-AP-

**Figure 1.**
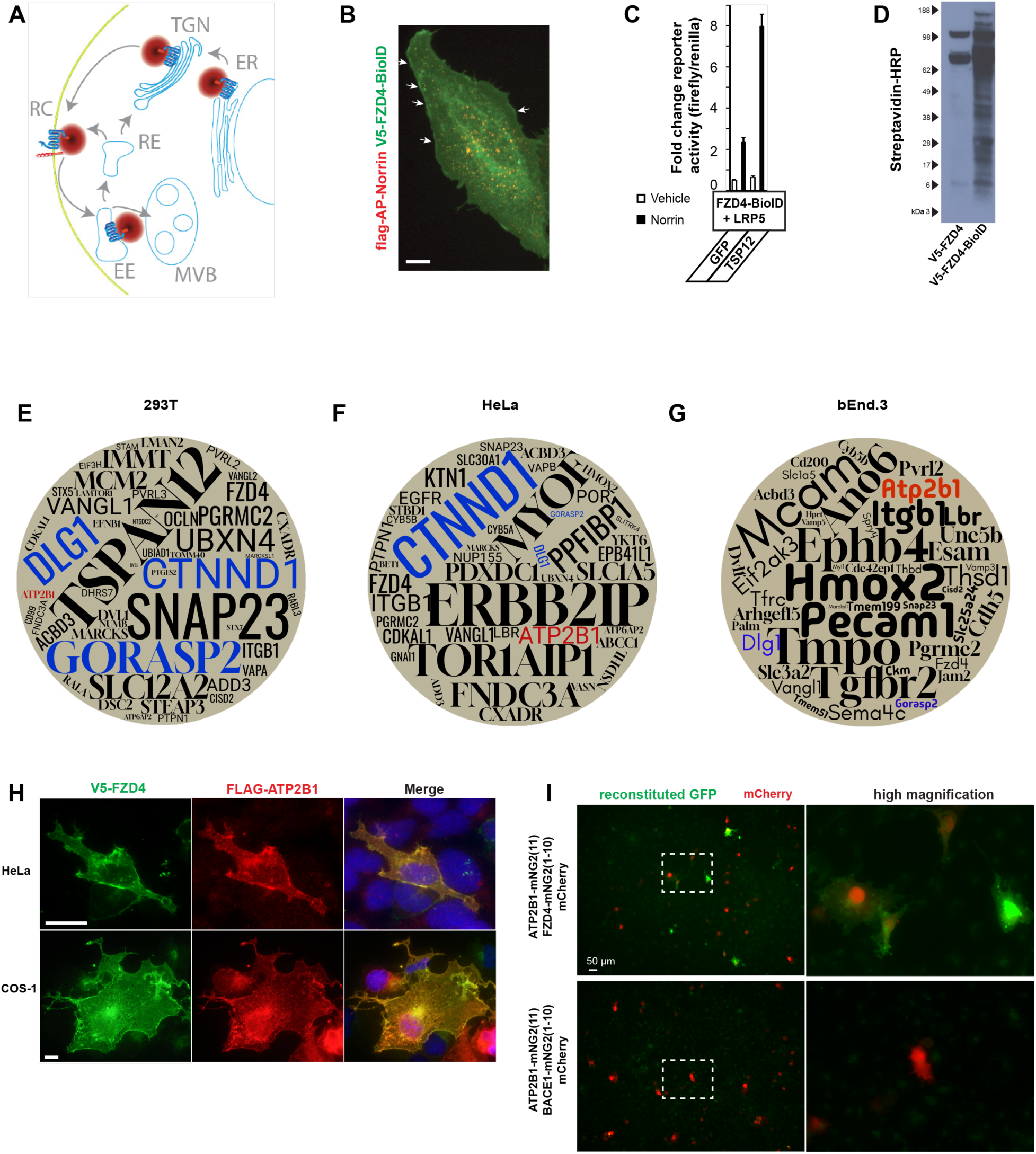
Identification of a proximity interaction of FZD4 and ATP2B1. **(A**) FZD4 with C-terminal BioID tag, schematic trafficking steps. Red cloud symbolizes 10 nm radius in which biotinylation is most likely to occur (not to scale). ER, endoplasmic reticulum, TGN, trans-Golgi network. RC, receptor complex. EE, early endosome. RE, recycling endosome, MVB, multi vesicular body. (**B**) FZD4-BioID reaches the plasma membrane (white arrows) and undergoes Norrin-induced endocytosis (intracellular puncta), as described for untagged FZD4. Representative of two experiments with similar results. Scale bar: 10 µm. (**C**) TOPFlash reporter assay shows activity of fusion protein in response to Norrin, mean +/- SD shown, n=3 biological replicates. (**D**) Streptavidin-HRP detects numerous biotinylated proteins in FZD4-BioID-transfected 293T cells, indicating that the BirA (R118G) component of the fusion protein is active. Representative of two experiments with similar results. (**E-G**) Filtering of LC-MS/MS data by specificity (vs. GFP-BioID), abundance, coverage and MS/MS count criteria yields prioritized lists of proximity interactors that are represented as word cloud, where the size of the letters correlates with the rank of the protein in the filtered list, see also Supplementary Fig. S2-4. Proteins of interest are highlighted in color. (**H**) Immuno-staining of V5-FZD4 and FLAG-ATP2B1 in HeLa and COS-1 cells. Scale bars: 10 µm. (**I**) Split-GFP reconstitution in COS-1 cells co-transfected with mCherry. The membrane protein BACE1 (β-secretase 1) was used as negative control.

Norrin binding to FZD4 and endocytosis, V5-FZD4-BioID was detected both at the plasma membrane as well as in endocytic vesicles (Fig. 1B), where it co-localized with flag-AP-Norrin (an efficiently secreted, soluble and bioactive alkaline-phosphatase fusion protein). This observation indicated that the FZD4 fusion protein underwent similar trafficking processes as FZD4 (without BioID tag) ^58^. TOPFlash reporter assays for Norrin-induced β-catenin-dependent FZD4 signaling in HEK 293T cells confirmed that the V5-FZD4-BioID fusion-protein transduced the Norrin signal. Signaling amplitude was increased in the presence of the Norrin co-receptor, TSPAN12, as expected ^12, 15, 59^, indicating that V5-FZD4-BioID initiated signaling and was able to cooperate with TSPAN12 (Fig. 1C). Immunoblots of 293T cell lysates transfected with V5-FZD5-BioID and treated with biotin showed substantial biotinylation of proteins of various molecular weights, whereas in lysates of V5-FZD4 expressing cells only few endogenously biotinylated proteins were detected with Streptavidin-HRP (Fig. 1D), confirming biotin ligase activity in the fusion protein. Next, we performed BioID screens in three cell lines that have been previously used in cell-based studies of Norrin/Frizzled4 signaling: HEK293T cells, HeLa cells, and bEnd.3 mouse brain endothelial cells. Collectively, these cell lines are suitable to identify mechanisms that regulate FZD4 in multiple biological contexts, including the BBB. In 293T cells, plasmids encoding LRP5 and TSPAN12 were co-transfected with the BioID constructs, whereas HeLa cells were transfected only with the BioID constructs, and bEnd.3 cells were virally transduced with vectors encoding GFP-BioID or V5-FZD4-BioID. Stable cell populations were generated by selecting blasticidin-resistant transduced cells. Staining for the V5 peptide tag showed FZD4 accumulated in the perinuclear compartment and at the plasma membrane in bEnd.3 cells (Supplementary Fig. S1). Biotinylated proteins were isolated from 293T, HeLa, and bEnd.3 cells and subjected to LC-MS/MS analysis. The bEnd.3 cell samples were processed through a different LC-MS/MS and bioinformatic pipeline than the 293T and HeLa cells (see methods). We filtered the proteins identified by LC-MS/MS for a combination of specificity in comparison with GFP-BioID, abundance, and protein coverage (Supplementary Fig. S2-S4), and displayed the shortlists of filtered proteins in a word cloud, in which the size of the letters correlates with semi-quantitative protein metrics (Fig. 1 E-G). A Venn diagram was generated to highlight that multiple proteins were identified in all three experiments, including FZD4 as an auto-proximity interaction, Discs Large homolog 1 (DLG1) and Golgi Reassembly-Stacking Protein 2 (GORASP2) (Supplementary Fig. S5). The robust identification of these proteins validated the screens, as DLG1 is a physical and functional FZD4 interactor ^60, 61^, and GORASP2 is a Golgi protein that controls FZD4 trafficking ^62^.

Additional proteins that were identified in all three cell lines include ATP2B1/PMCA1, a plasma membrane Ca^2+^-pump. Co-transfected TSPAN12, a Frizzled4 receptor complex component, was identified in 293T cells as expected ^15, 16^. Dishevelled-1 (DVL-1), which is an intracellular interactor of FZD4 ^58, 63^, further validated the screen in 293T cells.

Additional proteins of interest are addressed in the discussion. For further analysis, we focused on ATP2B1 because its function in ECs remains incompletely understood (see Introduction), its proximity interaction was unexpected, and siRNA-mediated knockdown of *Atp2b1* affected Norrin/Frizzled4 signaling more consistently than knockdown of other candidates (see below). To validate the spatial relationship between ATP2B1 and FZD4, we performed co-immunostaining of V5-FZD4 and FLAG-ATP2B1 in both HeLa and COS-1 cells (large cells suitable for co-localization studies), revealing co-localization at the plasma membrane (Fig. 1H). Furthermore, FLAG-ATP2B1-mNG2(11) reconstituted GFP fluorescence by association with co-transfected V5-FZD4-mNG2(1-10), whereas split GFP was not reconstituted with a BACE1-(aka β-secretase)-mNG2(1-10) control (Fig. 1 I). Together, we identified ATP2B1 as a proximity interactor of FZD4.

### ATP2B1 facilitates Norrin/Frizzled4 signaling in CNS ECs

To investigate a potential functional role of ATP2B1 in Norrin/Frizzled4 signaling, we employed three distinct siRNAs targeting different regions of *Atp2b1* (coding sequences and 3’UTR), of which *Atp2b1* siRNA #2 achieved only moderate knockdown efficiency in bEnd.3 cells, whereas *Atp2b1* siRNA #1 and #3 knocked down ATP2B1 efficiently on the mRNA (Fig. 2A) and protein level (Supplementary Fig. S6). We examined the expression of *Axin2*, an established target gene of the canonical Norrin/Frizzled4 signaling pathway in ECs ^34, 64^. In control siRNA-treated bEnd.3 cells, Norrin stimulation induced *Axin2* mRNA expression > 30-fold. As expected, this induction was abolished in FZD4-knockdown cells due to the absence of the receptor for Norrin/Frizzled4 signaling. Notably, *Atp2b1* knockdown, using each of three siRNAs, significantly attenuated Norrin-induced *Axin2* expression compared to controls, suggesting that ATP2B1 facilitates the Norrin/Frizzled4 pathway in ECs (Fig. 2B). RNAseq data of bEnd.3 cells revealed that there was no major expression of other ATP2B family paralogs (Supplementary Fig. S7). We used *P2ry*1 as another Norrin signaling target gene, which we identified from our bEnd.3 cells RNAseq data. *P2ry1* mRNA expression was induced by Norrin in control cells but not in FZD4-knockdown cells (Fig. 2C). *Atp2b1* knockdown reduced Norrin-induced *P2ry1* expression, consistent with the reduced expression of *Axin2*. To further corroborate these findings, we used our bEnd.3 cells stably expressing a TOPFlash reporter and Renilla luciferase ^65^ and observed that *Atp2b1* knockdown significantly reduced Norrin-induced TOPFlash activity compared to controls (Fig. 2D).

**Figure 2.**
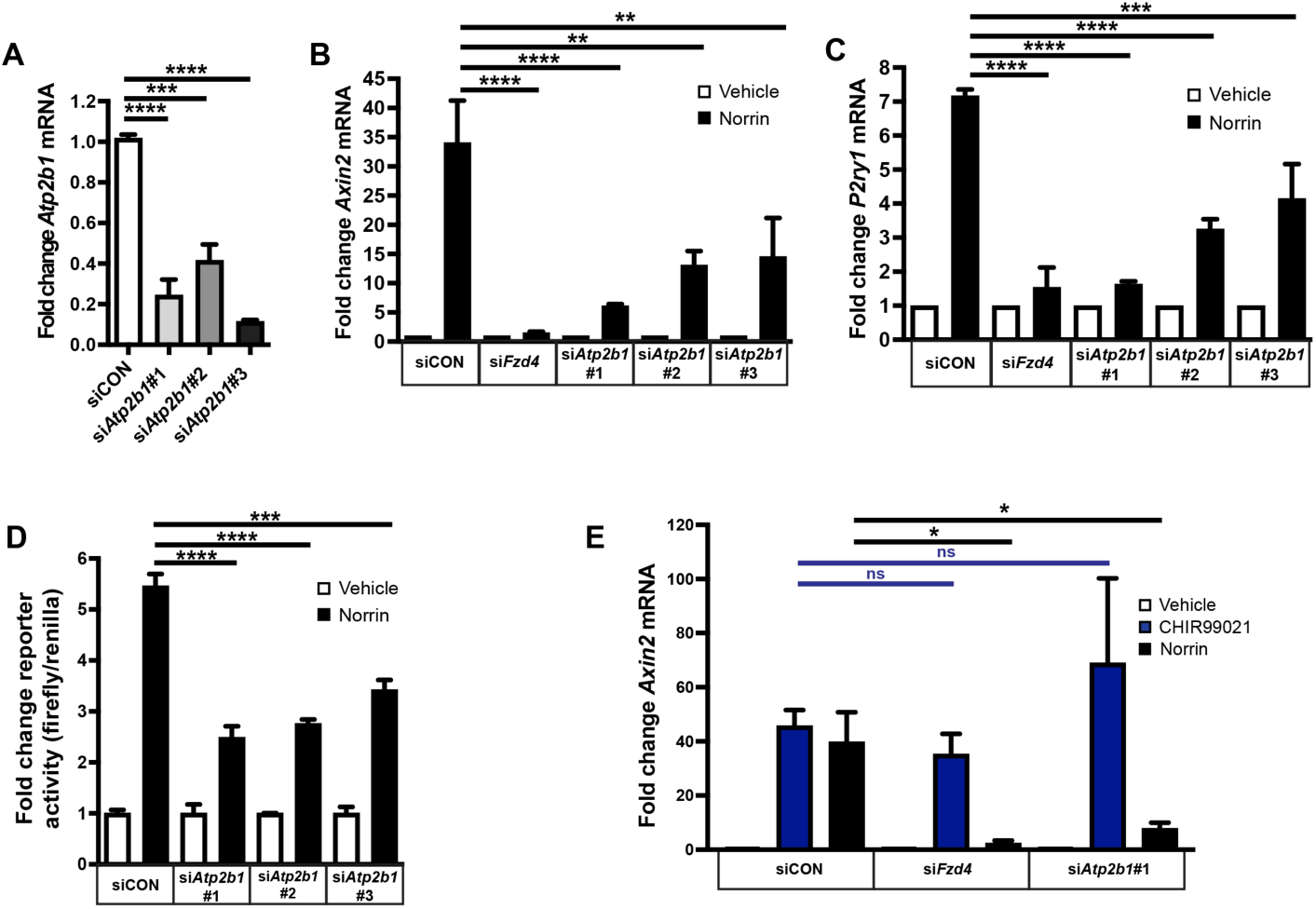
ATP2B1 facilitates Norrin/Frizzled4 signaling in CNS ECs. (**A**) Knockdown efficiency of ATP2B1 siRNAs in bEnd.3 cells assessed by RT-qPCR for *Atp2b1* mRNA. Two technical replicates per biological replicate were averaged, n=3 biological replicates, graphs represent mean +/- SEM, one-way ANOVA with Dunnett’s post hoc. In this and all following figures: *; P<0.05, **; P<0.01, ***; P<0.001 and ****; P<0.0001. (**B**) RT-qPCR of *Axin2*, a target gene of the canonical signaling, in bEnd.3 cells transfected with the indicated siRNAs and stimulated with vehicle (PBS, 0.1% BSA) or 200 ng/ml recombinant Norrin. Three independent experiments were performed, each with one biological replicate per condition. For each experiment, two technical replicates were averaged. The resulting values were subjected to paired, within-experiment normalization, in which the control condition was set to 1 and experimental conditions expressed relative to their matched control. n=3 biological replicates, graphs represent mean +/- SEM, one-way ANOVA with Dunnett’s post hoc. (**C**) RT-qPCR of *P2ry1* in bEnd.3 cells transfected with the indicated siRNAs and stimulated with vehicle or Norrin. *P2ry1* is an alternative target gene of canonical signaling based on our RNA sequencing in bEnd.3 cells (see Supplementary Fig. S7). Data were normalized and combined as described for panel B. Two technical replicates per biological replicate were averaged, n=3 biological replicates, graphs represent mean +/- SEM, one-way ANOVA with Dunnett’s post hoc. (**D**) Luciferase reporter assay using stable bEnd.3 TOPFlash cells. For each siRNA condition the data are normalized to the vehicle group to show the fold change of TOPFlash/renilla activity in response to Norrin. n=3 biological replicates, graphs represent mean +/- SEM, one-way ANOVA with Tukey’s post hoc. (**E**) RT-qPCR of *Axin2* mRNA after stimulation with Norrin or 2 µM of CHIR 99021, a GSK3β inhibitor. Data were normalized and combined as described for panel B. Two technical replicates per biological replicate were averaged, n=3 biological replicates, graphs represent mean +/- SEM, one-way ANOVA with Dunnett’s post hoc (blue bar; siCON+CHIR99021 as a comparison control, black bar; siCON+Norrin as a comparison control).

To determine the level at which ATP2B1 intersects with Norrin/Frizzled4 signaling, we utilized CHIR99021, a GSK3β inhibitor. GSK3β, a component of the β-catenin destruction complex, phosphorylates β-catenin, promoting its degradation ^66^. GSK3β inhibition activates β-catenin-dependent signaling independently of ligand-receptor interactions. In control cells, both Norrin and CHIR99021 robustly induced *Axin2* expression. *Atp2b1*-knockdown cells maintained their ability to upregulate *Axin2* expression in response to CHIR99021, but Norrin-induced activation was significantly impaired compared to controls (Fig. 2E). This differential response suggests that ATP2B1 deficiency impinges, at least in part, upstream of β-catenin, for example on the level of the receptor complex or signalosome. Using lentiviral transduction, we generated a stable bEnd.3 cell population expressing V5-FZD4 and performed a cell-surface biotinylation experiment. We found that *Atp2b1*-knockdown did not significantly affect FZD4 steady state levels at the cell surface, whereas Brefeldin A impaired cell-surface trafficking as expected (Supplementary Figure S8).

### Endothelial cell-specific *Atp2b1* gene ablation causes retinal angiogenesis and BRB phenotypes with variable expressivity

We re-analyzed our previously reported single-cell RNAseq of wild type mouse retina (Supplementary Fig. S9A) ^35^ and found broad *Atp2b1* mRNA expression across multiple retinal cell types, including vascular cells (Supplementary Fig. S9B). Additionally, the paralog *Atp2b4* was expressed in a more restricted manner, including in retinal vascular cells (Supplementary Fig. S9C). Next, we investigated the physiological role of *Atp2b1* in inducible endothelial cell-specific *Atp2b1* knockout (ECKO) mice, using a Cdh5-CreERT2 driver. The murine retinal vasculature develops in three distinct layers in a central to peripheral direction: the superficial vascular plexus in the nerve fiber layer (NFL) reaches the periphery by ∼P8, the deep plexus in the outer plexiform layer (OPL) reaches the periphery around P12, and the intermediate plexus in the inner plexiform layer (IPL) develops with slight delay and reaches the periphery by ∼P15 ^67^. Deep vascular plexus development is particularly sensitive to loss of Norrin signaling ^8^. We administered tamoxifen daily from P2-4 and analyzed the retinal vasculature at P13 to examine deep layer angiogenesis. The genotype of *Atp2b1* ECKO animals was *Atp2b1*^flox/-^; Cre^+^. Control genotypes were *Atp2b1*^flox/+^; Cre^-^, *Atp2b1*^flox/flox^; Cre^-^, *Atp2b1*^flox/-^; Cre^-^, or *Atp2b1*^flox/+^; Cre^+^, using littermates of the *Atp2b1* ECKO mice as control. All control mice exhibited normal vascular development of the superficial and intraretinal vasculature. In contrast, *Atp2b1* ECKO mice displayed vascular hypovascularization, particularly at the front of the superficial vascular plexus and in the deep vascular plexus (Fig. 3A). Incomplete deep vascular plexus development was occasionally concentrated to the OPL above major arteries. The expressivity of vascular phenotypes, which were present in all *Atp2b1* ECKO mice, was variable, with phenotypes ranging from only partial hypovascularization of the deep vascular plexus to virtually complete absence of the deep vascular plexus, glomeruloid vascular malformations, insufficiency of superficial vascular plexus development and invasion of hyaloid vessels into the retina. In retinas with mild phenotypes, 3D-analysis of the three retinal vascular layers revealed hypovascularization in the intraretinal capillary beds (Fig. 3B). These findings identified novel roles of ATP2B1 in retinal vascular ECs.

**Figure 3.**
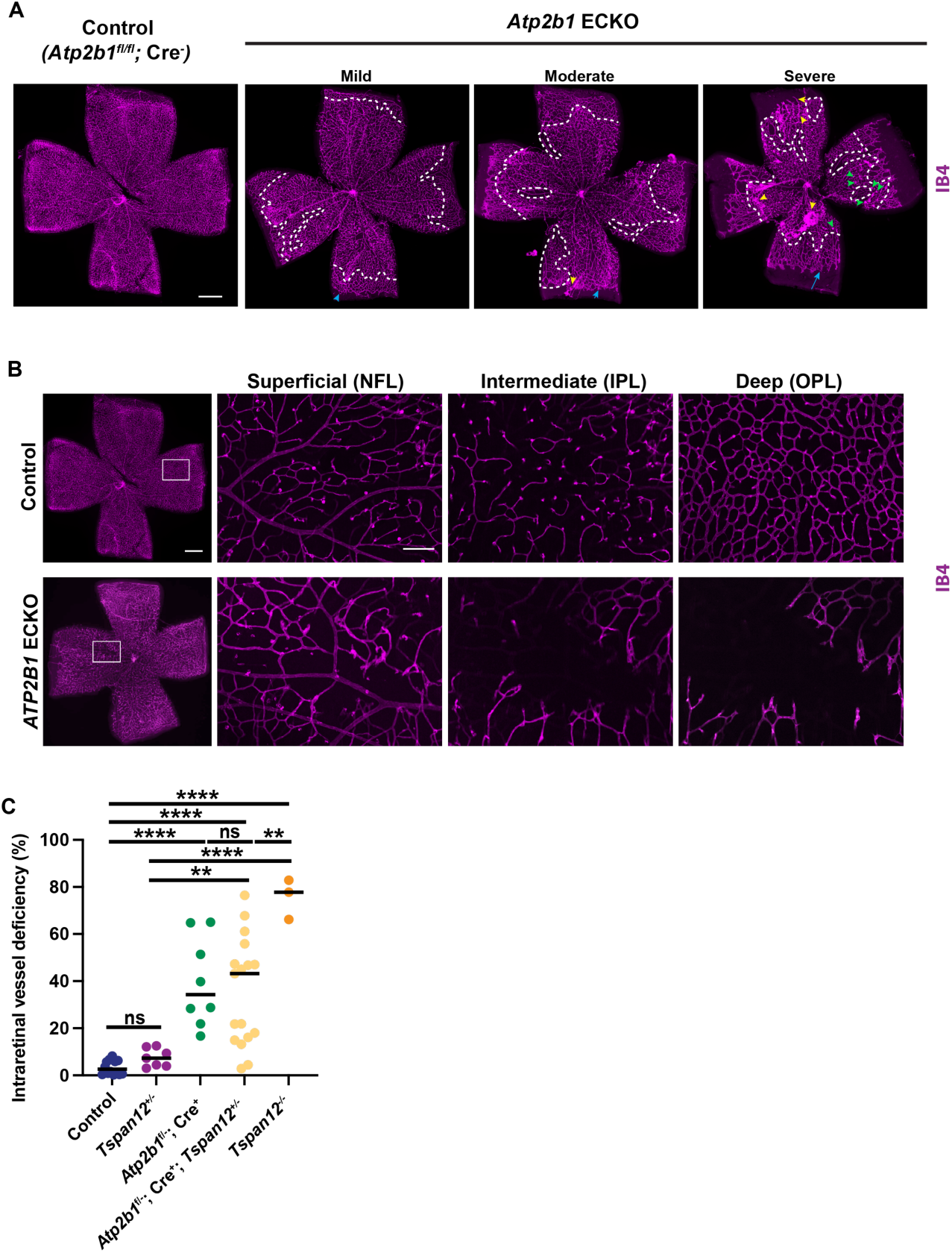
Vascular phenotypes in *Atp2b1* ECKO retinas. (**A**) *Atp2b1* ECKOs were induced with tamoxifen at P2, P3 and P4 and the retinas were harvested at P13. All *Atp2b1* ECKO retinas show a similar vascular phenotype, but the expressivity of the phenotype was variable and is illustrated in three examples. Blue arrows indicate a delay of the progression of the vascular front in the superficial vascular plexus. White dashed lines demarcate a boundary between areas with deep vascular plexus development vs. areas lacking the deep vascular plexus. Green arrowheads show glomeruloid vascular malformations. Yellow arrowheads show areas with anastomosis of retinal and hyaloid vasculature. Scale bar: 500 µm. (**B**) Left panels: Whole mount retinas of the indicated genotype are represented by stitched images. The white box indicates an area that is shown enlarged at 3 depth levels in the right panels. Scale bars: 500 µm left panel, 100 µm right panel. (**C**) Quantification of the area with intraretinal hypovascularization in an allele series of the indicated genotypes. n=3-14 retinas from 3-14 mice, graphs represent median, one-way ANOVA with Tukey’s post hoc.

Furthermore, the data suggested the possibility that our mouse colony on a C57BL/6J background may harbor unknown genetic modifiers that strongly influence the severity of the phenotype. Whether this modification represented an attenuation, or an exacerbation, remained unclear (see discussion) and variable phenotypes were observed in both sexes.

Quantification of intraretinal vessel deficiency confirmed the hypovascularization phenotype in the comparison of *Atp2b1* ECKO vs. control (Fig. 3C). To corroborate that ATP2B1 functions in the Norrin/Frizzled4 pathway, we attempted to test for genetic interactions of *Atp2b1* and the Norrin co-receptor *Tspan12* ^16, 17^. In the analysis of the allele series (Fig. 3C), we observed a non-significant trend towards an exacerbated hypovascularization in *Atp2b1* ECKO; *Tspan12*^+/-^ compound mutant retinas compared to *Atp2b1* ECKO retinas. This analysis was complicated by the high variability of the *Atp2b1* ECKO phenotype as well as the statistical power required for a comparison of 5 groups.

Because Norrin/Frizzled4 signaling is a key pathway for the induction and maintenance of the BRB ^9^, we investigated if BRB function was compromised. We assessed Plasmalemma Vesicle Associated Protein (PLVAP) expression, an EC fenestration/diaphragm component that is normally absent from retinal ECs, but is expressed in ECs of *Ndp*, *Lrp5*, and *Tspan12* gene ablated mice ^15^. While control retinas showed no substantial PLVAP expression (Fig. 4A), several *Atp2b1* ECKO mice exhibited localized PLVAP expression corresponding to regions of defective deep layer angiogenesis (Fig. 4A). The expressivity of the phenotype was highly variable, but the mean PLVAP^+^ area was significantly different between *Atp2b1* ECKO retinas and control retinas (Fig. 4B). Several *Atp2b1* ECKO; *Tspan12*^+/-^ mice displayed widespread PLVAP expression across the retinal vasculature, however, mean PLVAP expression in *Atp2b1* ECKO; *Tspan12*^+/-^ mice was similar to those in *Atp2b1* ECKO mice. We further used sulfo-NHS-LC-biotin, a small molecule biotinylation reagent that penetrates disrupted barriers ^68^ to evaluate BRB function. The increase in biotin^+^ area in *Atp2b1* ECKO retinas did not reach significance compared to controls, whereas leakage was significantly different in *Atp2b1* ECKO; *Tspan12*^+/-^ mice compared to controls (Fig. 4C).

**Figure 4.**
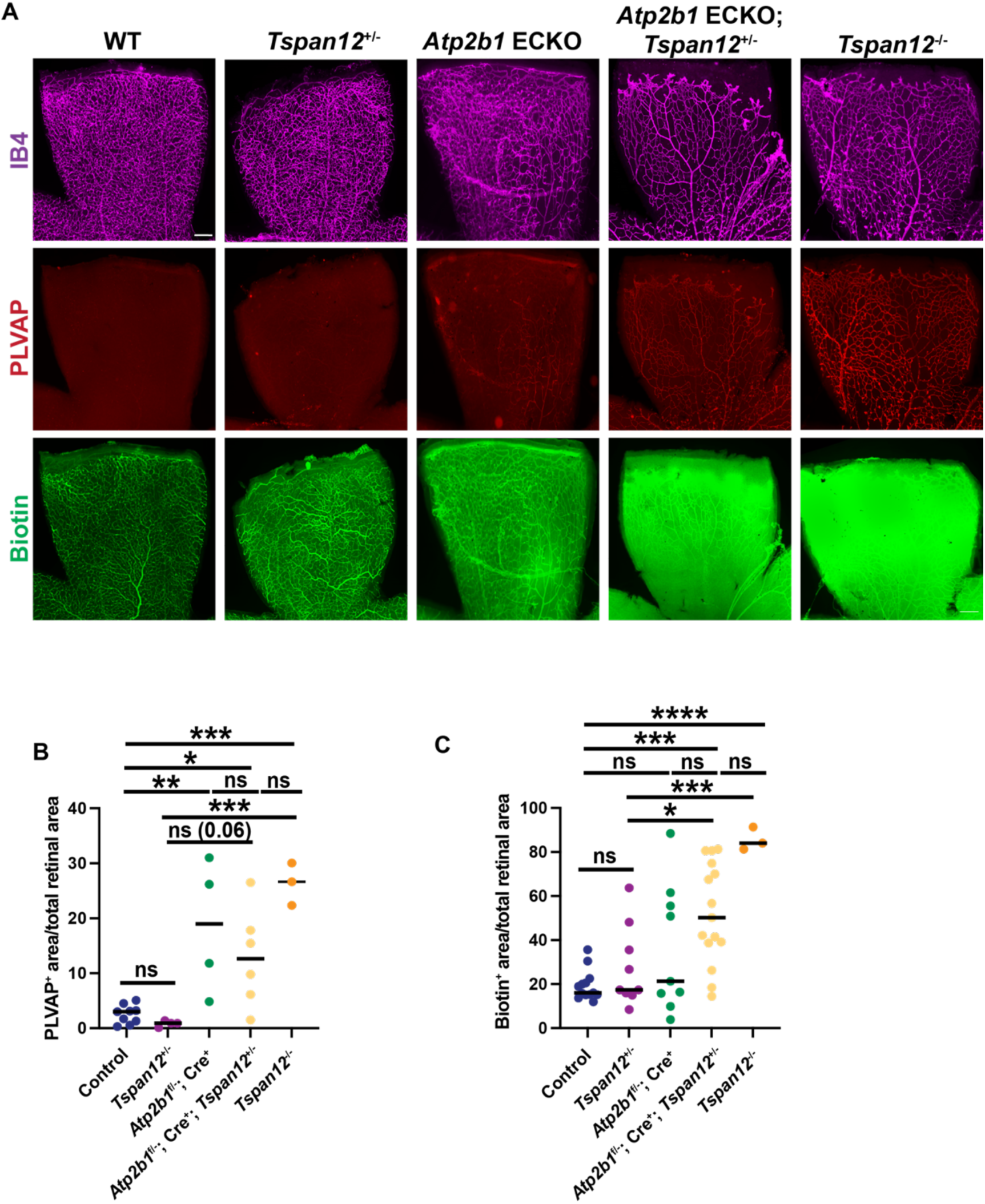
Analysis of BRB function in *Atp2b1* ECKOs. (**A**) Example images of retinal petals stained for IB4, PLVAP (a fenestration marker), and biotin (using streptavidin-Alexa 488) from an allele series with the genotypes indicated in the figure. Scale bar 200 µm. (**B**) Quantification of PLVAP^+^ area. n=3-9 retinas from 3-9 mice, graphs represent median, one-way ANOVA with Tukey’s post hoc. (**C**) Quantification of biotin^+^ area. n=3-15 retinas from 3-15 mice, graphs represent median, one-way ANOVA with Tukey’s post hoc.

In addition to the *Atp2b1* ECKOs of the *Atp2b1*^f/-^; Cre^+^ genotype, we also generated *Atp2b1* ECKOs with the genotype *Atp2b1*^f/f^; Cre^+^. The outcomes for hypovascuarization, PLVAP expression, and biotin leakiness were similar as in *Atp2b1*^f/-^; Cre^+^ retinas, albeit milder (Supplementary Fig. S10 A-C). We also analyzed aggregated data from *Atp2b1*^f/-^; Cre^+^ and *Atp2b1* ^f/f^; Cre^+^ ECKO retinas combined. In this analysis all phenotypic parameters (hypovascularization, PLVAP expression, biotin leakage) trended higher in the *Atp2b1* ECKO; Cre^+^; *Tspan12*^+/-^ group compared to *Atp2b1* ECKO, but the difference did not reach significance (Supplementary Fig. S10 D-F).

Together, these results demonstrate that EC-specific deletion of *Atp2b1* causes vascular defects and that the expressivity of the vascular phenotypes is variable (see discussion).

### Elevated intracellular Ca^2+^ inhibits Norrin/Frizzled4 signaling

Having established a role for ATP2B1 in Norrin/Frizzled4 signaling, retinal angiogenesis, and BRB maintenance, we investigated the mechanistic basis of this regulation. Given the function of ATP2B1 as a plasma membrane calcium pump, we hypothesized that its deletion would elevate intracellular calcium levels in bEnd.3 cells, especially since ATP2B1 is the major paralog in bEnd.3 cells (Supplementary Fig. S7). To test this hypothesis, we knocked down *Atp2b1* in bEnd.3 cells, which resulted in a significant increase of Fluo-4 signal (a membrane-permeable Ca^2+^ indicator dye whose emission was normalized to Hoechst fluorescence) compared to bEnd.3 cells transfected with non-targeting control siRNA (Fig. 5A and B). Next, we employed ionomycin, a calcium ionophore, to increase intracellular calcium concentrations in 293T cells co-transfected with FZD4, LRP5, TSPAN12 and the reporter constructs. In control cells, Norrin treatment induced a robust increase in luciferase activity. However, this response was dose-dependently and significantly attenuated by ionomycin (Fig. 5C). To further validate the calcium-dependent regulation of Norrin/Frizzled4 signaling, we utilized SEA0400, a selective inhibitor of the Na⁺-Ca²⁺ exchanger (NCX), which extrudes Ca^2+^ in its forward mode. 1 µM SEA0400 treatment of bEnd.3 cells suppressed Norrin-induced TOPFlash activity compared to controls (Fig. 5D). Consistent with these findings, ionomycin treatment significantly decreased the mRNA expression of Norrin target genes *Axin2* and *P2ry1* in bEnd.3 cells (Fig. 5, E and F). Collectively, these data support that elevated intracellular calcium, resulting from ATP2B1 deficiency, negatively regulates Norrin/Frizzled4 signaling in CNS ECs.

**Figure 5.**
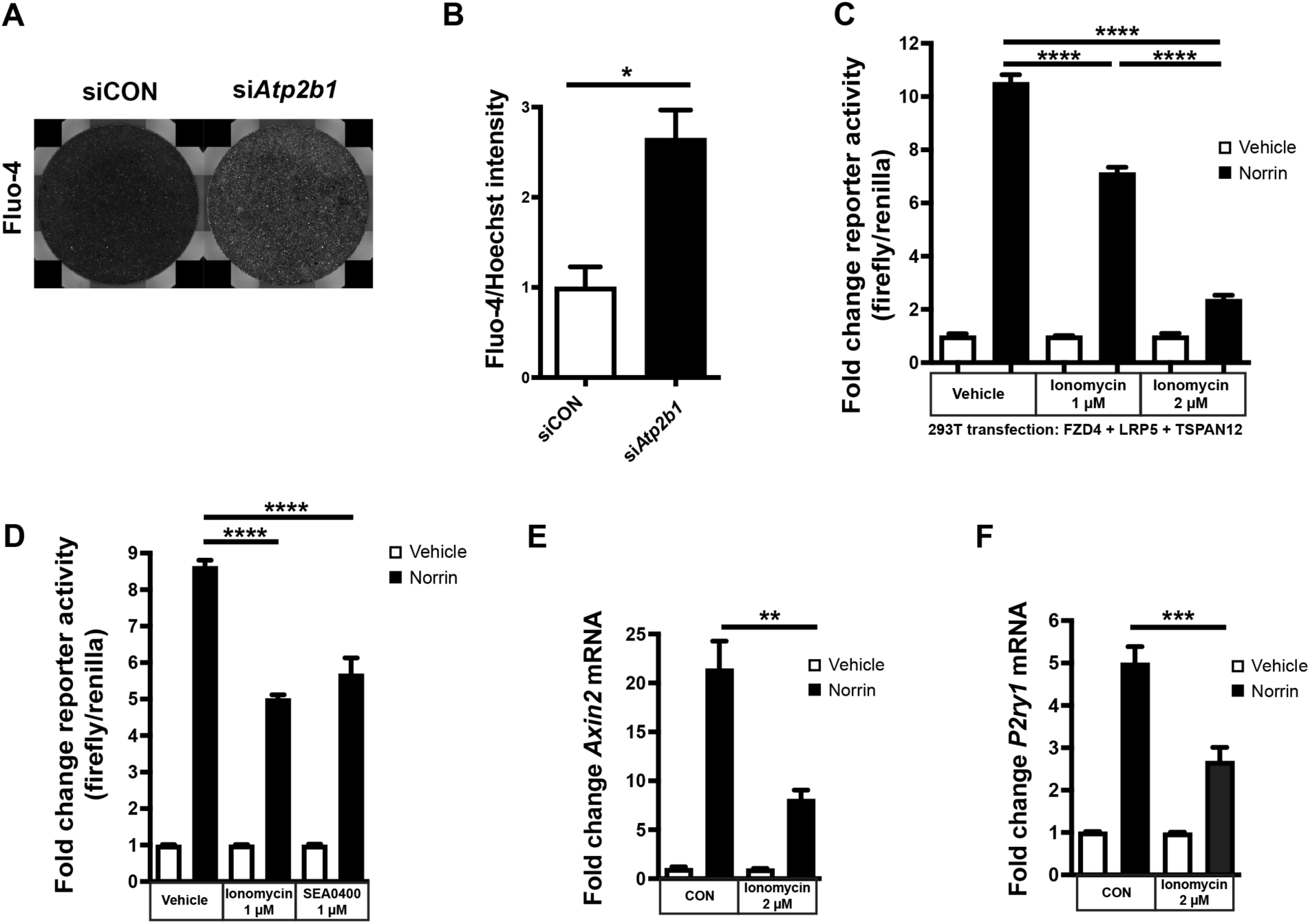
Elevated intracellular Ca^2+^ inhibits Norrin/Frizzled4 signaling. **(A**) Images of two wells of a 48-well plate containing bEnd.3 cells loaded with Fluo4 Ca^2+^ indicator. Cells were transfected with siRNAs as indicated in the figure. (**B**) Quantification of Fluo-4 intensity normalized by Hoechst intensity. n=3 biological replicates, graphs represent mean +/- SEM, unpaired Student’s t-test. (**C**) 293T cells transfected with the indicated constructs and TOPFlash and CMV-Renilla plasmids were stimulated with vehicle or Norrin in the presence or absence of ionomycin. n=3 biological replicates, graphs represent mean +/- SEM, one-way ANOVA with Tukey’s post hoc. (**D**) Stable bEnd.3 TOPFlash cells stimulated with vehicle or Norrin in the presence of ionomycin or SEA0400, a Na⁺-Ca²⁺ exchanger inhibitor. n=3 biological replicates, graphs represent mean +/- SEM, one-way ANOVA with Tukey’s post hoc. (**E**) Total RNA from bEnd.3 cells subjected to RT-qPCR for *Axin2* mRNA after stimulation with vehicle or Norrin in the presence of DMSO or ionomycin. Two technical replicates per biological replicate were averaged, n=3 biological replicates, graphs represent mean +/- SEM, one-way ANOVA with Tukey’s post hoc. (**F**) Total RNA from bEnd.3 cells subjected to RT-qPCR for *P2ry1* mRNA after stimulation with vehicle or Norrin in the presence of DMSO or ionomycin. Two technical replicates per biological replicate were averaged, n=3 biological replicates, graphs represent mean +/- SEM, one-way ANOVA with Tukey’s post hoc.

### Calcineurin-mediated NFAT activation negatively regulates Norrin/Frizzled4 signaling

Because several Nuclear Factor of Activated T cells (NFAT) family transcription factors (NFATc1-4) are controlled by the Ca^2+^-dependent phosphatase calcineurin, we investigated whether NFAT-signaling modulates Norrin/Frizzled4 signaling. Intracellular calcium promotes calmodulin-dependent activation of calcineurin, which dephosphorylates multiple NFAT paralogs, facilitating their nuclear translocation and subsequent target gene transcription ^69^. Our RNA sequencing of bEnd.3 cells showed that NFAT2 (aka NFATc1), NFAT4 (aka NFATc3) and Ca^2+^-independent NFAT5 are the predominant family members in bEnd.3 cells (Supplementary Fig. S7). We assessed the expression of known NFAT2 target genes, *Egr3* and *Nr4a2* ^70^, in bEnd.3 cells treated with *Atp2b1* or non-targeting control siRNA (Fig. 6A and B). ATP2B1 knockdown significantly induced the expression of both NFAT2 target genes compared to controls. Treatment with the calcineurin inhibitor cyclosporin A (CsA), which suppresses NFAT activity, significantly reduced *Egr3* and *Nr4a2* expression in *Atp2b1*-knockdown cells, confirming that these target genes are under control of calcineurin/NFAT signaling in ECs. Furthermore, we transfected EGFP-hNFAT2 with C-terminal EE tag into bEnd.3 cells treated with non-targeting siRNA or *Atp2b1* siRNA. While NFAT2 localized predominantly outside the nucleus in control cells, we observed significant translocation to the nucleus after *Atp2b1* knockdown (Fig. 6C and D). To investigate the relationship of NFAT signaling and Norrin/Frizzled4 signaling, we inhibited NFAT signaling in bEnd.3 TOPFlash cells. CsA treatment rescued the reduction of canonical Norrin/Frizzled4 signaling after *Atp2b1* knockdown (Fig. 6E). This finding indicates that calcineurin/NFAT activation contributes to inhibiting the Norrin/Frizzled4 pathway. Furthermore, we used lentivirus to transfer constitutively active NFAT2 (aka NFATc1) into bEnd.3 cells and selected a stable population using blasticidin. This population exhibited significantly elevated expression of *Egr3* and *Nr4a2* compared to parental cells (Fig. 6F and G).

**Figure 6.**
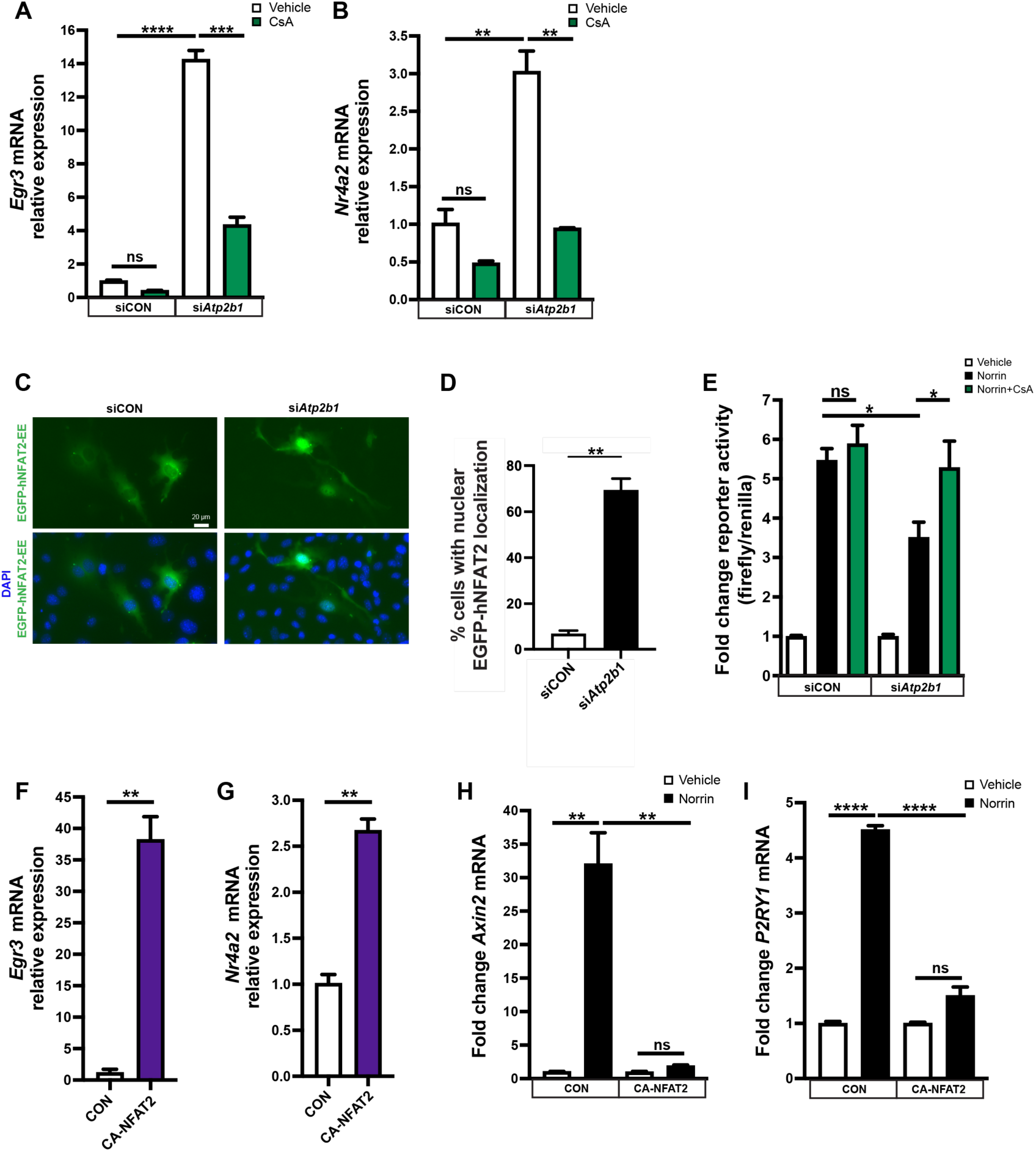
ATP2B1 deficiency regulates Norrin/Frizzled4 signaling through the calcineurin/NFAT pathway. (**A-B**) bEnd.3 cells were transfected with *Atp2b1* siRNA#3 or control siRNA and analyzed with RT-qPCR for *Erg3 or Nr4a2* after stimulation with vehicle 1 µg/ml cyclosporin A, a calcineurin inhibitor. Two technical replicates per biological replicate were averaged, n=2 biological replicates, graphs represent mean +/-SEM, one-way ANOVA with Tukey’s post hoc. (**C-D**) bEnd.3 cells transfected with EGFP-hNFAT2-EE and the indicated siRNAs were scored for nuclear NFAT2 localization to determine NFAT2 activation. The EE tag is glutamate-rich tag derived from the polyoma virus medium T antigen. 3 images per biological replicate were averaged, n=3 biological replicates, graphs represent mean +/- SD, homoscedastic t-test. (**E**) Stable bEnd.3 TOPFlash cells were transfected with the indicated siRNAs and stimulated with vehicle, Norrin or Norrin and 1 µg/ml cyclosporin A. n=9 biological replicates, graphs represent mean +/- SEM, one-way ANOVA with Tukey’s post hoc. (**F-G**) bEnd.3 cells were infected with CA-NFAT2 lentivirus and selected with blasticidin. mRNA expression of *Erg3* and *Nr4a2* was assessed by RT-qPCR in parental bEnd.3 cells vs. CA-NFAT2 expressing cells. Two technical replicates per biological replicate were averaged, n=2 biological replicates, graphs represent mean +/- SEM, unpaired t-test. (**H-I**) Parental or CA-NFAT2 expressing bEnd.3 cells were stimulated with vehicle or Norrin and *Axin2* and *P2ry1* mRNA expression was assessed by RT-qPCR. Two technical replicates per biological replicate were averaged, n=2 biological replicates, graphs represent mean +/- SEM, one-way ANOVA with Tukey’s post hoc.

While Norrin treatment successfully induced the expression of its target genes *Axin2* and *P2ry1* in control bEnd.3 cells, this response was virtually abolished in CA-NFAT2-overexpressing cells (Fig. 6H and I). Together, these results support that ATP2B1 deficiency activates NFAT signaling, which in turn contributes to suppressing Norrin/Frizzled4 signaling.

### ATP2B1 facilitates WNT7A/B signaling and brain angiogenesis

Similar to Norrin/Frizzled4 signaling in retinal vascular development, neuroepithelial-derived WNT7A and WNT7B orchestrate critical aspects of neurovascular development in the developing brain and spinal cord. These ligands activate β-catenin-dependent signaling in ECs, thereby promoting both angiogenesis and BBB specialization. Next to Frizzled receptors, GPR124 functions as an essential co-activator in the WNT7A/B signaling axis (see introduction). To investigate the role of ATP2B1 in Wnt/β-catenin signaling, we used 293T cells transfected with plasmids encoding WNT7A or WNT7B and co-cultured these cells with siRNA-treated bEnd.3 cells carrying the TOPFlash reporter. While control siRNA-treated cells activated canonical signaling in response to WNT7A and WNT7B, signaling amplitude was significantly reduced in *Atp2b1* knockdown cells (Fig. 7A). To further elucidate the requirement of ATP2B1 in Wnt/β-catenin signaling *in vivo*, we generated *Atp2b1 ECKO; Gpr124*^-/-^ compound mutant mice (Cre^-^ littermates were used as control, this included littermates that were heterozygous for the *Gpr124* null or *the Atp2b1* floxed allele). After maternal intraperitoneal injection of tamoxifen at E10.5, E11.5, and E12.5, brain sections were collected from embryonic day 13.5 (E13.5) embryos (Fig. 7B). Previous studies have established that *Gpr124* knockout embryos display pronounced defects in vascularization of the cerebral cortex and medial ganglionic eminence (MGE), accompanied by EC de-specialization and reduction of glucose transporter GLUT1 expression ^14^. Immunohistochemical analysis revealed that both wild-type and *Atp2b1* ECKO mice exhibited largely normal vascularization patterns in the cerebral cortex and MGE. In contrast, both *Gpr124* KO and *Atp2b1* ECKO; GPR124^-/-^ compound mutant mice displayed glomeruloid vascular structures in the cerebral cortex and aberrant vascular aggregation in the MGE (Fig. 7C). Notably, the severity of these phenotypes was significantly more pronounced in the *Atp2b1* ECKO; *Gpr124*^-/-^ mice compared to *Gpr124* single KO mice, indicating that ATP2B1 deficiency exacerbates the angiogenesis defects observed in *Gpr124*^-/-^ mice (Fig. 7D and E). Collectively, these data support that ATP2B1 genetically interacts with components of WNT7A/B signaling during brain vascular development.

**Figure 7.**
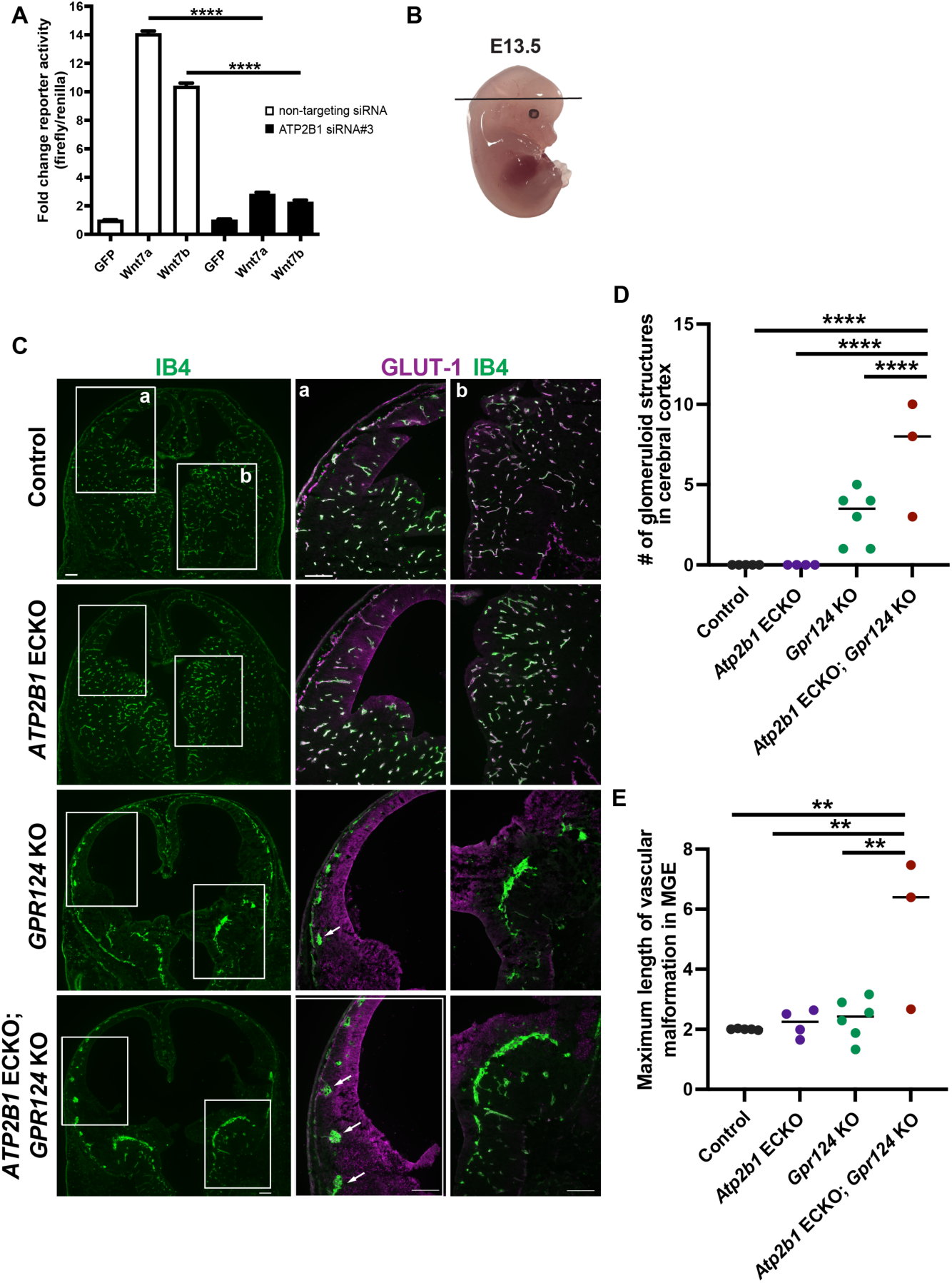
ATP2B1 regulates WNT7A/B-induced Frizzled signaling in the developing brain. (**A**) Stable bEnd.3 TOPFlash cells were co-seeded with 293T cells, the latter were transiently transfected with plasmids encoding WNT7A or WNT7B before co-seeding. n=3 biological replicates, graphs represent mean +/- SEM, one-way ANOVA with Tukey’s post hoc. (**B**) Orientation of sections analyzed in panel C. (**C**) E13.5 forebrain 20 µm sections of the indicated genotype. White boxes show areas in the two developing hemispheres and are shown enlarged in the right panels. White boxes labeled a show the developing forebrain cortex, white boxes labeled b show the medial ganglionic eminence (MGE). White arrows point to glomeruloid vascular malformations. Scale bars: 200 µm. (**D**) Quantification of a total number of glomeruloid vascular malformations in the cerebral cortex. n=3-6 embryos, graphs represent median, one-way ANOVA with Tukey’s post hoc. (**E**) Maximum length of vascular malformation in MGE, n=3-6 biological replicates, graphs represent median, one-way ANOVA with Tukey’s post hoc.

## DISCUSSION

A major conclusion from this study is that ATP2B1 facilitates Norrin- and WNT7A/B-signaling in CNS ECs. This conclusion is supported by the effects of *Atp2b1* reduction in multiple independent cell-based signaling assays and the phenotypes of genetic mouse models. Mechanistically, increased Ca^2+^/NFAT signaling associated with *Atp2b1* knockdown is involved in suppressing Norrin/Frizzled4 signaling. The mechanisms downstream of NFAT remain to be determined, but it is noted that NFAT can exert its functions via both transcriptional ^70^ and non-transcriptional mechanisms ^71^. The evidence that ionomycin or CA-NFAT2 overexpression each reduce Norrin/Frizzled4 signaling does not rule out that ATP2B1 deficiency could affect Norrin/Frizzled4 signaling via additional mechanisms besides NFAT signaling. *Atp2b1* knockdown in ECs did not inhibit canonical signaling induced by the GSK3β inhibitor CHIR99021 to the same extent as Norrin/Frizzled4 signaling, suggesting functions of ATP2B1 or Ca^2+^/NFAT signaling upstream of β-catenin.

A second major conclusion is that ATP2B1 functions in the regulation of retinal angiogenesis and the BRB. There is overlap between phenotypes resulting from loss of Norrin/Frizzled4 signaling and *Atp2b1* LOF in the retinal vasculature. In both situations deep vascular plexus formation is impaired, the fenestration marker PLVAP is upregulated, and BRB dysfunction is evident ^72^. Yet, given the broad roles of Ca^2+^ in EC signal transduction ^48, 52, 53^, *Atp2b1* ECKO phenotypes may not be exclusively due to impaired Norrin/Frizzled4 signaling. Norrin and WNT7A/B signaling are pathways that enable angiogenesis in the CNS but are not required for systemic angiogenesis. We observed angiogenesis phenotypes in *Atp2b1* ECKO mice in the retina, and exacerbated angiogenesis phenotypes in *Gpr124*;*Atp2b1* ECKO compound mutant mice in the embryonic brain. Further studies are needed to determine if ATP2B1 functions in ECs are CNS-specific.

Our finding that Ca^2+^/NFAT signaling inhibits Norrin/Frizzled4 signaling implies that factors that regulate EC Ca^2+^, including VEGF ^53^, ROS ^73^, thrombin ^48, 74^, bradykinin ^75, 76^ or mechanosensory stress ^51^ could reduce the levels of Norrin/Frizzled4 signaling in ECs and contribute to increased vascular permeability, for example in diabetic retinopathy. The data suggest that other genes that are involved in Ca^2+^ homeostasis, including the ATP2B1 paralog ATP2B4, may functionally interact with β-catenin-dependent signaling in ECs. Furthermore, Ca^2+^-signaling and Wnt/β-catenin signaling potentially intersect in additional biological contexts ^77^.

Our study revealed additional proximity interactions of FZD4 beyond ATP2B1 (Supplemental Fig. S5). These interactions include DLG1, GORASP2, CTNND1/p120, and ITGB1. DLG-1 is a PDZ domain scaffold protein that was identified as a Frizzled4 interactor in a yeast-two-hybrid screen ^61^. Importantly, EC-specific gene ablation of *Dlg1* results in a reduction of retinal deep vascular plexus density and increased PLVAP expression. Furthermore, *Dlg1* genetically interacts with *Fzd4* and *Tspan12* with respect to retinal vascular phenotypes, and with *Tspan12* and *Ndp* regarding BBB phenotypes. These phenotypes are rescued by stabilized β-catenin. DLG1 interacts with FZD4 via PDZ domains 1 and 2 ^60^. This study provides compelling evidence for complex formation of DLG1 and FZD4 and our identification of DLG1 among the proximity interactors in three cell lines validates the proximity biotinylation screens on a technical level. GORASP2 promotes the transport of cargo with PDZ recognition motifs, including FZD4 ^62^. Thus, our screen identified at least two proteins that can interact with the PDZ recognition motif of FZD4. Additional proteins related to trafficking that were identified in all three cell lines include ACBD3 (a Golgi-resident protein) and SNAP23 (a T-SNARE). Another protein consistently identified in all three cell lines is ITGB1. The significance of this finding is unclear, but it is noted that the ITGB1 interactor integrin-linked kinase (ILK) is implicated in regulating retinal angiogenesis through β-catenin-dependent signaling and is genetically linked to FEVR ^78^. Furthermore, we identified Catenin Delta 1 (aka p120-catenin, CTNND1) in 293T and HeLa cells. This is of interest, as a role for CTNND1 in β-catenin signaling has been attributed to binding LRP5/6 and the regulation of internalization of the signalosome into multi-vesicular bodies ^79^, which is thought to lead to β-catenin stabilization ^58, 80^. Mutations in human *CTNND1* or mouse *Ctnnd1* cause retinal hypovascularization reminiscent of FEVR, due to roles in adherens junctions, or β-catenin-dependent signaling, or both ^81^. The finding of a proximity interaction between CTNND1 and FZD4 suggests that additional roles of CTNND1 in FZD4-signaling remain to be uncovered.

FZD4 proximity interactors may have functions beyond β-catenin-dependent signaling, for example, the identification of VANGL1 in all three cell lines stands out. Furthermore, there may be functionally significant interactors that we detected only in one of three cell lines, e.g., some EC-specific interactors may not be expressed in 293T or HeLa cells.

UNC5B was only detected in bEnd.3 ECs, and this protein is implicated in BBB function^82^. TSPAN12, which is physically associated with FZD4 ^16^, was co-transfected into 293T cells and was only identified in those cells. Finally, proximity interactors may not necessarily modulate FZD4 function, instead, FZD4 may modulate the function of the nearby proteins. Together, the proximity biotinylation screens yielded numerous interactors of interest and led to the identification of ATP2B1 as a facilitator of Norrin and WNT7A/B signaling. Our mechanistic studies identify Ca^2+^/NFAT as an inhibitor of β-catenin-dependent signaling in ECs and imply that pathways and pathological conditions that increase EC Ca^2+^ have the potential to modulate β-catenin-dependent signaling and regulate angiogenesis and barrier function. In addition, this mechanism may be relevant in the context of pharmacological approaches that transiently disrupt endothelial barriers for drug delivery.

### Limitations of the study

Our genetic experiments conducted on a C57BL/6J background revealed significant variability in the retinal vascular phenotypes of *Atp2b1* ECKO mice and the reason(s) for this variability remain undefined. Possible explanations include variable efficiency of recombination, variable expression of redundant factors in ECs (e.g., ATP2B4), or variable compensation in Ca^2+^-handling. The fact that the machinery regulating intracellular Ca^2+^ is highly complex may make *Atp2b1* ECKO mice susceptible to the effect of genetic modifiers acquired through genetic drift.

Whether *Atp2b1* ECKO mice compounded with homozygous loss of *Tspan12*. *Tspan12* KO retinas display a genetic interaction remains to be determined. The complete lack of deep vascular plexus formation and very severe BRB dysfunction in *Tspan12* KO mice may obscure the detection of further reduction of deep vascular plexus formation and BRB function when ATP2B1 is reduced in addition to TSPAN12.

Our identification of proximity interaction, co-localization, and split-GFP reconstitution is consistent with proximity of FZD4 and ATP2B1, but experimental support for complex formation or the identification of a direct interaction interface is currently lacking. The mechanisms by which NFAT signaling intersects with Norrin/Frizzled4 signaling in ECs remain to be investigated.

## RESOURCE AVAILABILITY

### Lead contact

Further information and requests for resources and reagents should be directed to and will be fulfilled by the lead contact Harald Junge.

### Materials availability

Materials generated in the Junge laboratory will be available from the Lead Contact, but may require a completed Materials Transfer Agreement (MTA).

### Data and code availability

- RNA-seq data have been deposited at NCBI GEO under accession number GSE299490.
- Proteomics data are available in the PRIDE database under PXD065058 and PXD077093
- This study does not report original code.
- Any additional information required to reanalyze the data reported in this paper is available from the lead contact upon request.

## Acknowledgements

We thank Heidi Roehrich, and Danielle May for excellent technical support. We thank the Center for Metabolomics and Proteomics at the University of Minnesota for generation of quantitative proteomics data and the Viral Vector and Cloning Core at the University of Minnesota for generating lentivirus. This study was supported by grants from the NIH (R01EY024261 and R01EY033316 to HJJ and 1R21DA056728-01A1 to ZC) and the Wallin Neuroscience Discovery Award (to HJJ). NIH grant P30GM145398 for the Pediatric Research COBRE to KJR supported the Biochemistry Core who performed BioID pulldowns from HeLa and 293T cells. JL was supported by T32EY025187. EK was supported by T32-NS109604. HJJ has been named to the Mary and Marian Robinson Chair in Macular Degeneration.

## Author contributions

HJJ, HNJ, and CZ designed experiments. HNJ, EK, LZ, CZ, EO, QCD, ZC, JL, MH and HJJ conducted experiments and analyzed data. KJR assisted with proteomics workflows, KDF provided the *Atp2b1* floxed mouse strain reagents. HNJ and HJJ wrote the manuscript. All authors edited the manuscript.

## Declaration of generative AI and AI-assisted technologies in the writing process

During the preparation of this work the authors used ChatGPT to edit the manuscript draft. After using this tool, the authors reviewed and edited the content as needed and take full responsibility for the content of the publication.

## SUPPLEMENTAL INFORMATION

Document S1. Figures S1-S10 and supplemental methods.

## STAR* METHODS

## 1. EXPERIMENTAL MODEL AND STUDY PARTICIPANT DETAILS

### Animals

*Atp2b1* floxed mice (*Atp2b1^tm1c(KOMP)Wtsi^)* were generated from *Atp2b1^tm1a(KOMP)Wtsi^* sperm by Dr. Klaus-Dieter Fischer’s lab ^47^ and imported to Minnesota with permission from MMRRC UC Davis. *Atp2b1* floxed mice were imported on a C57BL/6J background and subsequently backcrossed at the Minnesota site for 7 additional generations using C57BL/6J mice obtained from Jackson laboratories. Experimental mice were up to 13 days old (P13). The *Atp2b1* null allele was generated by recombining the floxed *Atp2b1* allele in the germline using Sox2-Cre (Jackson lab stock # 008454). Tg(Cdh5-cre/ERT2)1Rha ^83^ mice were used for EC-specific recombination and provided by Dr.

Ralf Adams under MTA with CANCERTOOLS.ORG. The *Tem5* aka *Gpr124* null allele ^18^ was kindly provided by Dr. Brad St. Croix under MTA with NCI. Studies were conducted using animals of both sexes and data were not analyzed disaggregated for sex as the study containing multiple compound mutant genotypes was not powered for this analysis. Mice were housed in a specific pathogen-free animal facility. The Animal Care and Use Committee of the University of Minnesota, Twin Cities, approved all animal protocols.

### Cell lines

bEnd.3 cells, COS-1 cells, HEK293T cells, and HeLa cells (all female-derived) were cultured in high-glucose DMEM supplemented with 10% FBS at 37 °C in a humidified atmosphere containing 5% CO₂.

## 2. METHOD DETAILS

### Plasmids and virus

pcDNA 3.3 V5-hFZD4-BioID was generated by PCR-cloning the BioID coding sequence into the BamHI and AgeI sites of pcDNA3.3 V5-hFZD4-DmrAA, which we described previously ^16^. This construct was used for transient transfections into 293T and HeLa cells. pLenti-CMV-V5-hFZD4-BioID-IRES-BlastR and pLenti-CMV-V5-hFZD4 were generated by PCR amplification and subcloning into the AgeI and XhoI sites of a pLenti-CMV-MCS-IRES-BlastR vector. This vector was obtained by excising the AgeI/XhoI turboRFP insert from Addgene plasmid #102343 (deposited by Ghassan Mouneimne) and replacing it with a primer dimer harboring a MCS. pLenti-CMV-flag-hATP2B1-IRES-BlastR was generated by subcloning a gene synthesis-derived flag-hATP2B1 fragment into the BamHI and NheI sites of pLenti-CMV-MCS-IRES-BlastR – this fragment harbors silent mutations that transfer resistance to specific siRNAs. pLenti-CMV-3xHA-CA-NFAT2-IRES-BlastR was generated by PCR cloning 3xHA-CA-NFAT2 into the AgeI and XhoI sites of pLenti-CMV-MCS-IRES-BlastR. The 3xHA-CA-NFAT2 sequence was obtained from Addgene plasmid #11102, deposited by Anjana Rao. pcDNA3.3 hWnt7b was described previously ^16^. pcDNA Wnt7a was obtained from Addgene (#35914, deposited by Marian Waterman). Virus was generated by the UMM Viral Vector and Cloning Core.

### TOPFlash luciferase assay

293T were transfected with a plasmid mix that we described previously ^16^, in brief: 4 ng pcDNA.3.3 hFZD4, 8 ng pcDNA.3.3 hLRP5, 8 ng pcDNA3.3 hTSPAN12 and 140 ng reporter mixture were transfected with TransIT LTI according to manufacturer’s instruction per 48 well. 6-8 hours after transfection, cells were stimulated with 200 ng/ml Norrin (R+D Systems, 3014-NR-025) for 12-16 hours and processed with the Promega Dual-Glo lysis buffer and substrates according to manufacturer’s instructions. For raising intracellular Ca^2+^, ionomycin was present from the time cells had adhered and maintained throughout the experiment. For TOPFlash experiments in bEnd.3 cells, we used our previously described ^65^ stable bEnd.3 population infected with two lentivirus encoding TOPFlash (puromycin resistance) and CMV-Renilla (blasticidin resistance), cultured under selection pressure in high glucose DMEM, 10 % FBS, 1 % penicillin/streptomycin, 1 μg/ml puromycin and 10 μg/ml blasticidin. 30K cells were seeded per 48-well in full DMEM without puromycin and blasticidin and incubated overnight. Medium was replaced (200 µl per well) and cells were stimulated for 18 h with 200 ng/ml Norrin (R+D Systems) before harvest with Dual Glo lysis buffer and substrates (Promega, E1910). For raising intracellular Ca^2+^, ionomycin or SEA0400 (or DMSO as vehicle) was present 16 hours before adding norrin and maintained throughout the experiment. For bEnd.3 TOPFlash experiments in the presence of Cyclosporin A, cells were transfected with siRNAs. 48 hours after transfection, cells were stimulated for 24 hours with vehicle, Norrin, or Norrin in the presence of 1 µg/mL CsA. TOPFlash data are normalized to the respective condition without stimulus (-Norrin) within one experimental group, where each siRNA transfection is a separate experimental group. To stimulate bEnd.3 TF cells with WNT7A or WNT7b, bEnd.3 TF cells were seeded into a 6 well plate (30K cells per well) and transfected the following day with either 20 nM final concentration of control siRNA or ATP2B1 siRNA #3 and incubated for 48 hours. 293T cells were seeded into a 6 well plate (1 million cells per well) and transfected 6 hours later with 2 µg pcDNA3.3 EGFP, pcDNA-WNT7a, or pcDNA3.3 hWnt7b) expression vectors. Transfected 293T cells were incubated for 42 hours post-transfection. For co-culture, siRNA-transfected bEnd.3 TOPFlash cells (30K cells per well) were combined with GFP- or Wnt7- transfected 293T cells (60K cells per well) in a 48 well plate. After 24 hours of co-culture, cells were subjected to the Dual Glo luciferase assay.

### Norrin-induced FZD4 endocytosis

HeLa cells were transfected with FZD4 and subjected to endocytosis as described previously ^58^. In brief: 36 hours after transfection, cells were incubated with ice cold conditioned medium containing flag-alkaline phosphatase-Norrin for one hour on ice, washed with cold medium, and transferred to 37°C for 1 hour to induce endocytosis. Non-internalized flag-AP-Norrin was removed by incubation with 150 mM NaCl, 50 mM glycine, pH 2.4 for 3 minutes at RT before cells were fixed, permeabilized, and stained using antibodies directed against the V5-tag and Flag-tag.

### BioID proximity labeling

293T, HeLa, or bEnd.3 cells were cultured as described above. For the experiment using 293T cells, four 10 cm diameter dishes per condition were each seeded with 6 million cells. 3 h later, each dish was transfected with 400 ng pcDNA.3.3 V5-hFZD4-BioID or GFP-BioID, 800 ng pcDNA.3.3 hLRP5, 800 ng pcDNA 3.3 hTSPAN12, 6000 ng empty vector, 2000 ng pcDNA.3.3 EGFP using 15 µL TransIT-LTI. After 24 h, medium was replaced and 50 µM biotin was added for 24 hours. 293T cells were harvested by scraping, washed in PBS twice, and frozen. Sample IDs were S479 and S482 for duplicate GFP-BioID samples, and S480 and S483 for duplicate FZD4-BioID samples. For the experiment using HeLa cells, 2 million cells were seeded into each of four 10 cm diameter dishes and transfected with 250 µl linear PEI per 10 µg total plasmid DNA (9 µg pcDNA3.3 EGFP, which we used as filler vector for vector balancing, plus 1 µg of either V5-FZD4-BioID or GFP-BioID). After 24 hours, cells were incubated for an additional 24 hours with 50 µM biotin. Cells were washed in PBS, harvested using enzyme free cell dissociation solution (Gibco #13151-014), and frozen. GFP-BioID was sample S455, V5-FZD4-BioID was sample S456. For the experiment using bEnd.3 cells, cells were infected with lentivirus for gene transfer of V5-FZD4-BioID or GFP-BioID. After selection with blasticidin, four 150 cm^2^ dishes per sample were incubated for 24 hours with 50 µM biotin. Duplicate samples were generated. Cells were washed in PBS and directly lysed. Cells from all experiments were processed according to a detailed method paper ^84^. In brief, cells were lysed in 8 M urea 50 mM Tris, pH 7.4, buffer with protease inhibitor (87785; Thermo Fisher Scientific) and DTT under sonication, precleared with gelatin-sepharose beads (17095601; Cytiva), incubated with streptavidin-sepharose beads (17511301; Cytiva/GE Healthcare), washed in 8 M urea, 50 mM Tris, pH 7.4, and resuspended in 50 mM ammonium bicarbonate saturated with 1 mM biotin.

### Protein digestion and mass spectrometry

Samples were reduce-alkylated in 8 M urea, 50 mM ammonium bicarbonate buffer with 10 mM tris(2-carboxyethyl)phosphine (TCEP) at 30 °C for 60 min and 30 mM iodoacetamide (IAA) in the dark at room temperature for 30 min. Samples were diluted to 1M urea and digested with mass spec grade Trypsin/LysC mix overnight (V5071; Promega). The peptide eluates were acidified with formic acid and purified via a C18 matrix (5190-6532; Agilent) using Agilent AssayMap BRAVO liquid handling system.

Solvent was removed in a SpeedVac and subjected to LC-MS/MS. The 293T and HeLa samples were analyzed at the SBP Discovery Core San Diego using Proxeon EASY nanoLC system coupled to a Q-Exactive Plus mass spectrometer (Thermo Fisher Scientific). MS spectra were analyzed with MaxQuant 1.5.5.1 software and searched against the Homo sapiens Uniprot protein sequences database (version January 2018) and GPM cRAP sequences (common contaminants). Precursor mass tolerance was set to 20 ppm for the initial mass recalibration and then 4.5 ppm for the main search.

Product ion mass tolerance was set at 0.5 Da and maximum precursor ion charge state was 7. Carbamidomethylation of cysteines was searched as a fixed modification, while oxidation of methionines and acetylation of protein N-terminal were searched as variable modifications. Enzyme was set to trypsin with max 2 missed cleavages. The target-decoy-based false discovery rate (FDR) filter for spectrum and protein identification was set to 1%. The bEnd.3 cells were processed at the University of Minnesota Center for Mass Spectrometry and Proteomics. Approximately 200 ng of reconstituted peptides were analyzed using an Orbitrap Fusion mass spectrometer (Thermo Fisher Scientific) coupled to a Thermo UltiMate 3000 RSLCnano system. Mass spectra were analyzed using SEQUEST within Proteome Discoverer 4.0. Additional details on LC-MS/MS acquisition and database searching are provided in the Supplement.

### Filtering of priority candidates

Candidate interactors were prioritized based on multiple criteria: an FZD4/GFP label-free quantification (LFQ) abundance ratio >10 for HeLa and HEK293T cells and >5 for bEnd.3 cells; sequence coverage thresholds of >20% (HeLa), >15% (HEK293T), and >14% (bEnd.3); inclusion among the top 50 genes by absolute LFQ abundance in FZD4-BioID samples; MS/MS spectral count filters (excluding proteins with <5 counts in HeLa and HEK293T or <3 counts in bEnd.3); and finally sorted by the absolute MS/MS count of FZD4-BioID. Selected candidate interactors from each dataset were visualized using WordCloud.com and are also displayed in the supplement in tabular form.

### Immunofluorescence colocalization assay

30K cells HeLa cells were seeded into a 8-well chamber slide and transfected with 80 ng pcDNA3.3 V5-FZD4 and 80 ng pLenti-CMV-flag-hATP2B1-IRES-BlastR using 0.3 µL TransIT LTI (Mirus). Twenty-four hours post-transfection, cells were fixed for 10 minutes with ice cold methanol, washed three times with PBS, and then blocked and permeabilized for 30 minutes in PBS containing 0.1% Triton X-100 and 5% goat serum. Cells were incubated overnight at 4 °C with the following primary antibodies: mouse α-V5 (1:500, Bio-Rad, MCA1360) and rabbit α-Flag (1:500, Cell Signaling clone D6W5B #14793). The next day, cells were washed three times with blocking buffer and incubated with secondary antibodies: goat α-mouse Alexa-488 (1:2000, Invitrogen, A11001), goat α-rabbit Alexa-555 (1:2000, Invitrogen, A21428), and DAPI (1:1000) for nuclear staining. After final washes, cells were mounted using Fluoromount-G (Invitrogen, 00-4958-02). Images were acquired using a Leica DMIL epifluorescent microscope or a Keyence BZ-X810 digital microscope.

### Split-GFP reconstitution

40K COS-1 cells were seeded into a gelatin-coated 8-well chamber-slide (LABTEK) and transfected with pLenti-CMV-flag-hATP2B1-mNG2(11) and pcDNA3.3 V5-hFZD4-mNG2(1-10) or BACE1-mNG2(1-10) and mCherry using a total of 160 ng plasmid DNA and 0.3 µl TransIT-LTI (Mirus). After 48 hours, cells were fixed with 4% PFA for 10 min at RT and imaged using a Keyence BZ-X810 digital microscope.

### siRNA knockdown in bEnd.3 cells

bEnd.3 cells were cultured as described above. 300K cells were seeded in each well of a 6-well plate and allowed to adhere overnight. The following day, cells were transfected with siRNA at a final concentration of 20 nM using Lipofectamine RNAiMAX (Invitrogen, 56532) according to the manufacturer’s instructions. For Western blot analysis of knockdown efficiency after 48 hours, bEnd.3 cell lysates were probed with anti-ATP2B1(Abcam ab190355) and anti-β-actin (Novus NB600-501H). For TOPFlash reporter experiments in bEnd.3 cells, 48 hours after siRNA transfection, cells were treated with either recombinant Norrin (R+D Systems, 200 ng/mL) or the GSK3β inhibitor CHIR99021 (2 µM) in fresh DMEM. Cells were incubated for an additional 24 hours before RNA extraction.

Mouse *Atp2b1* and *Fzd4* siRNA was from Sigma. Nucleotides are ribonucleotides unless otherwise denoted with d for deoxynucleotide. si*Atp2b1* #1 (5’-GUCAUGGGCCAGUGGUCAA-dTdT-3’ and 5’- UUGACCACUGGCCCAUGAC-dTdT-3’), si*Atp2b1* #2 (5’-GGCUAAACACGAUCUCUGU-dTdT -3’ and 5’-ACAGAGAUCGUGUUUAGCC-dTdT-3’), si*Atp2b1* #3 (5’-CUUUAUACCUCCUAAGAAG-dTdT-3’ and 5’-CUUCUUAGGAGGUAUAAAG-dTdT-3’) and si*Fzd4* (5’- CCUGUUAUUUCUAUGAAAU-dTdT-3’ and 5’-AUUUCAUAGAAAUAACAGG-dTdT-3’). AllStars Neg. Control siRNA (Qiagen, 1027281) was used as non-targeting control.

### RNA isolation and quantitative PCR

Total RNA was extracted from cultured cells using TRIzol reagent (APB Bioscience, FP312) according to the manufacturer’s instructions. Equal amounts of total RNA were reverse-transcribed into complementary DNA (cDNA) using the Maxima First Strand cDNA Synthesis Kit for RT-qPCR (Thermo Fisher Scientific, K-1642). Quantitative PCR (qPCR) was carried out using SYBR Green master mix, and relative gene expression levels were calculated using the comparative Ct (ΔΔCt) method. Primers were designed to span exon–exon junctions to avoid genomic DNA amplification. Primer sequences are provided in Table S1

### Bulk RNA-sequencing

bEnd.3 cells were stimulated with vehicle (PBS, 0.1% BSA) or 200 ng/ml Norrin (R+D systems, 3014-NR-025) for 24 hours. Total RNA was extracted from quadruplicate samples using Trizol (Invitrogen, Cat# 15596026) according to manufacturer’s instructions. Genomic DNA contamination was removed with TURBO DNA-free DNase treatment (Invitrogen, AM1907). Libraries were made using the TakaraBio SMARTer Pico Mammalian kit. Novaseq 150PE was carried out, generating approximately 45 million reads per sample. Sequence alignment to the mouse reference genome was conducted using the RNA-Seq module of SeqMan NGen (DNASTAR Lasergene).

Differential expression analysis was carried out using ArrayStar (DNASTAR Lasergene). Data are available at NCBI GEO under GSE299490.

### FZD4 cell-surface biotinylation

A stable population of V5-FZD4 expressing bEnd.3 cells was generated by lentiviral transduction and was maintained under blasticidin selection (10 µg/ml). 300K cells per 6-well were seeded into 2 ml of full DMEM without blasticidin. The next day, cells were transfected with 20 µM final siRNAs as described above. 48 hours later, cell-surface

biotinylation was performed. Brefeldin A treatment (3 µg/ml) was for 24 hours before cell surface biotinylation. Cultures were transferred to ice to prevent endocytosis, the medium was removed, and cells were briefly rinsed with ice-cold PBS (pH 8.0). Surface proteins were labeled by incubating intact cells with 1.5 ml PBS (pH 8.0) containing 0.25 mg/ml EZ-Link Sulfo-NHS-SS-biotin (Thermo Fisher) for 45 min on ice with gentle rocking. Excess reagent was removed and then biotinylation was stopped by adding 400 µl PBS supplemented with 200 mM glycine (pH 8.0) for 5 min at 4 °C. Cells were washed with 2 ml cold TBS. Cells were lysed in 0.8 ml cold RIPA buffer containing 0.5 mM CaCl₂, 2 mM MgCl₂ (to improve DNAseI activity), 0.2 mg/ml DNase I, and EDTA-free protease inhibitor cocktail. Lysis proceeded for 20 min on ice with intermittent vortexing, followed by addition of EDTA to 20 mM final. Insoluble material was removed by centrifugation (10 min, 20,000 × g, 4 °C). For input samples, 104 µl of cleared lysate was mixed with 16 µl 1M DTT and 40 µl 4× LDS sample buffer and incubated at 45 °C for 10 min. The remaining lysate was incubated with 50 µl pre-equilibrated NeutrAvidin agarose (Thermo Fisher) for 1 h at 4 °C with rotation to capture biotinylated proteins.

Beads were transferred to spin filter columns connected to a vacuum manifold and washed 10 × 0.8 ml with 0.1x RIPA diluted with 150 mM NaCl solution at room temperature. Bound proteins were eluted by incubation with 105 µl of 1x LDS sample buffer containing 100 mM DTT for 20 min at 45 °C with agitation, followed by collection of eluates by centrifugation. Samples were analyzed by immunoblotting using HRP-conjugated antibodies against V5 (Bio-Rad MCA1360P).

### Tamoxifen preparation and administration

Tamoxifen (Sigma, T5648) was dissolved in sterile corn oil (Sigma, C8267) by rotating overnight at RT while protected from light using aluminum foil. The resulting solution was sterile filtered, aliquoted, and stored at −80 °C for up to one month. For Cre induction, postnatal pups received 50 µg tamoxifen s.c. once daily on postnatal days P2, P3, and P4. Embryos received Tamoxifen through a pregnant dam by intraperitoneal injections of 100 µL tamoxifen solution (20 mg/mL; 2 mg per injection) once daily on embryonic days E10.5, E11.5 and E12.5.

### Vascular analysis of retinal whole mounts

Mice were anesthetized using an isoflurane drop jar and euthanized via cervical dislocation. Eyes were enucleated and fixed immediately in 4% PFA for 15 minutes at RT. Retinas were dissected and blocked in 5% goat serum with 0.5% Triton X-100 in PBS for 1 hour at RT. Immunostaining was performed overnight at 4 °C in blocking buffer with primary antibodies: mouse α-PLVAP (1:100, BD Bioscience, 550563) and secondary antibodies: Goat α-rat cross-adsorbed Alexa-555 (1:500, Invitrogen, A21434), Isolectin B4 Alexa-647 (1:100, Invitrogen, I32450), and α-streptavidin Alexa-488 (1:400, Invitrogen, S11223). Between staining steps, retinas were washed five times for 30 minutes each in PBS containing 0.1% Triton X-100 at RT with shaking. Retinas were post-fixed and mounted for imaging.

Images were acquired using a Keyence BZ-X810 digital microscope. Whole retina images were captured using 10× objective with tile stitching. Areas lacking intermediate or deep vascular plexuses were manually measured using ImageJ. Individual vascular layers were captured at 20× magnification. PLVAP^+^ area and biotin^+^ area were quantified from 10× stitched images using a threshold established from control samples, which was consistently applied across all genotypes using ImageJ. PLVAP^+^ area and biotin-^+^ area was normalized by total retina area.

### Administration of biotin

To assess vascular permeability, sulfo-NHS-LC-biotin (Thermo Fisher, 21335) was administered intraperitoneally at a dose of 10 µL/g body weight (20 mg/mL solution) to postnatal day 13 (P13) pups. The biotin tracer was allowed to circulate for 60 minutes prior to sacrifice. Eyes were enucleated, and retinas were immersion-fixed in 4% PFA for 15 minutes at RT. Biotinylated proteins were visualized using 1:400 streptavidin conjugated to Alexa Fluor 488 (Invitrogen, S11223).

### Fluo-4 assay

bEnd.3 cells were seeded into a 48 well plate at ∼50% confluency. The day after plating, the cells were transfected with a control siRNA or ATP2B1 siRNA #3. After 72 hours, intracellular calcium was measured using the Fluo-4 Direct Calcium Assay Kit (Thermo Fisher Scientific, F10472). A 2X Fluo-4 Direct calcium reagent loading solution was prepared following the manufacturer’s protocol and was supplemented with 5 mM probenecid and 2 µg/mL Hoechst 33342. An equal volume of the 2x loading solution was added to the cells (e.g., 250 µL/well for a 48 well plate, the final concentration for probenecid is 2.5 mM and for Hoechst 33342 is 1 µg/mL). After incubation at 37°C for 30 min to allow Fluo-4 and Hoechst 33342 dyes to load into the cells, the fluorescent signals from Fluo-4 and Hoechst 33342 were captured and quantified using a Celigo Image Cytometer (Revvity). For each well, the integrated Fluo-4 fluorescence intensity per well was normalized to that from Hoechst 33342.

### NFATC2 nuclear localization

35K bEnd.3 cells in full high-glucose DMEM were seeded in a gelatin-coated 8-well chamberslide (LABTEK) and transfected with 20 nM final siRNA 2 hours later. After additional 14 hours, the medium was replaced. Additional 2 hours later, cells were transfected with 50 ng EGFP-hNFAT2-EE-WT (Addgene #24219, deposited by Jerry Crabtree) plus 110 ng empty vector using 0.3 µl Lipofectamine 3000 and 0.3 µl P3000 condenser. After additional 36 hours, cells were fixed with 4% PFA for 10 min at RT and imaged using a Keyence BZ-X810 digital microscope. Cells were counted in two categories: nuclear GFP signal weaker than cytoplasm, and nuclear GFP signal stronger than cytoplasm.

### Immunostaining of embryonic brain sections

Embryos were collected at E13.5 and fixed in 4% PFA overnight. Subsequently, embryos were washed with PBS and immersed in 30% sucrose for 24 hours. Embryos were then rinsed in PBS and frozen in OCT. Embryonic brains were sectioned in 20 µm thickness and blocked in 5% goat serum with 0.5% Triton X-100 in PBS for 1 hour at RT. Brains were incubated overnight at 4 °C with the following primary antibodies: rabbit α-mouse Glut-1 (1:200, Cell signaling, 73015T). After washing, the following secondary antibody or lectin were applied: Goat α-rabbit Alexa-647 (1:1000, Invitrogen, A21245) and Isolectin B4 Alexa-488 (1:100, Invitrogen, I21411). Images were acquired using a Keyence BZ-X810 digital microscope.

## 3. QUANTIFICATION AND STATISTICAL ANALYSIS

Statistical analyses were conducted using either the Statistics Kingdom web application or GraphPad Prism software. The specific statistical tests used are indicated in the corresponding figure legends. The sample size (n) is provided and defined in the figure legends. Data were assessed for normality and variance and are presented as mean ± standard error of the mean (SEM) or standard deviation (SD), as indicated in the figure legends. For comparisons among multiple groups, one-way ANOVA followed by Tukey’s post hoc test or Dunnett’s post hoc were used for normally distributed data, while the Kruskal–Wallis test was applied for non-parametric data. Two-group comparisons were performed using an unpaired t-test, Welch’s t-test, or the Mann–Whitney U test, as determined by the Statistics Kingdom app. A p-value < 0.05 was considered statistically significant, with significance denoted as follows: *P < 0.05; **P < 0.01; ***P < 0.001; ****P < 0.0001.

## REFERENCES

1. Sweeney MD, Zhao Z, Montagne A, Nelson AR, Zlokovic BV. Blood-Brain Barrier: From Physiology to Disease and Back. Physiol Rev 99, 21–78 (2019).

2. Obermeier B, Daneman R, Ransohoff RM. Development, maintenance and disruption of the blood-brain barrier. Nat Med 19, 1584–1596 (2013).

3. Langen UH, Ayloo S, Gu C. Development and Cell Biology of the Blood-Brain Barrier. Annu Rev Cell Dev Biol 35, 591–613 (2019).

4. Diaz-Coranguez M, Ramos C, Antonetti DA. The inner blood-retinal barrier: Cellular basis and development. Vision Res 139, 123–137 (2017).

5. Liebner S, Dijkhuizen RM, Reiss Y, Plate KH, Agalliu D, Constantin G. Functional morphology of the blood-brain barrier in health and disease. Acta Neuropathol 135, 311–336 (2018).

6. Liebner S, et al. Wnt/beta-catenin signaling controls development of the blood-brain barrier. J Cell Biol 183, 409–417 (2008).

7. Xu Q, et al. Vascular development in the retina and inner ear: control by Norrin and Frizzled-4, a high-affinity ligand-receptor pair. Cell 116, 883–895 (2004).

8. Ye X, et al. Norrin, frizzled-4, and Lrp5 signaling in endothelial cells controls a genetic program for retinal vascularization. Cell 139, 285–298 (2009).

9. Wang Y, Rattner A, Zhou Y, Williams J, Smallwood PM, Nathans J. Norrin/Frizzled4 signaling in retinal vascular development and blood brain barrier plasticity. Cell 151, 1332–1344 (2012).

10. Daneman R, Agalliu D, Zhou L, Kuhnert F, Kuo CJ, Barres BA. Wnt/beta-catenin signaling is required for CNS, but not non-CNS, angiogenesis. Proc Natl Acad Sci U S A 106, 641–646 (2009).

11. Stenman JM, Rajagopal J, Carroll TJ, Ishibashi M, McMahon J, McMahon AP. Canonical Wnt signaling regulates organ-specific assembly and differentiation of CNS vasculature. Science 322, 1247–1250 (2008).

12. Wang Y, et al. Interplay of the Norrin and Wnt7a/Wnt7b signaling systems in blood-brain barrier and blood-retina barrier development and maintenance. Proc Natl Acad Sci U S A 115, E11827–E11836 (2018).

13. Smallwood PM, Williams J, Xu Q, Leahy DJ, Nathans J. Mutational analysis of Norrin-Frizzled4 recognition. J Biol Chem 282, 4057–4068 (2007).

14. Zhou Y, Nathans J. Gpr124 controls CNS angiogenesis and blood-brain barrier integrity by promoting ligand-specific canonical wnt signaling. Dev Cell 31, 248–256 (2014).

15. Junge HJ, et al. TSPAN12 regulates retinal vascular development by promoting Norrin- but not Wnt-induced FZD4/beta-catenin signaling. Cell 139, 299–311 (2009).

16. Lai MB, et al. TSPAN12 Is a Norrin Co-receptor that Amplifies Frizzled4 Ligand Selectivity and Signaling. Cell Rep 19, 2809–2822 (2017).

17. Bruguera ES, Mahoney JP, Weis WI. The co-receptor Tetraspanin12 directly captures Norrin to promote ligand-specific beta-catenin signaling. Elife 13, (2025).

18. Cullen M, et al. GPR124, an orphan G protein-coupled receptor, is required for CNS-specific vascularization and establishment of the blood-brain barrier. Proc Natl Acad Sci U S A 108, 5759–5764 (2011).

19. Kuhnert F, et al. Essential regulation of CNS angiogenesis by the orphan G protein-coupled receptor GPR124. Science 330, 985–989 (2010).

20. Vanhollebeke B, et al. Tip cell-specific requirement for an atypical Gpr124- and Reck-dependent Wnt/beta-catenin pathway during brain angiogenesis. Elife 4, (2015).

21. America M, Bostaille N, Eubelen M, Martin M, Stainier DYR, Vanhollebeke B. An integrated model for Gpr124 function in Wnt7a/b signaling among vertebrates. Cell Rep 39, 110902 (2022).

22. Eubelen M, et al. A molecular mechanism for Wnt ligand-specific signaling. Science 361, (2018).

23. Posokhova E, et al. GPR124 Functions as a WNT7-Specific Coactivator of Canonical beta-Catenin Signaling. Cell Rep 10, 123–130 (2015).

24. Yuki K, et al. GPR124 regulates murine brain embryonic angiogenesis and BBB formation by an intracellular domain-independent mechanism. Development 151, (2024).

25. Cho C, Smallwood PM, Nathans J. Reck and Gpr124 Are Essential Receptor Cofactors for Wnt7a/Wnt7b-Specific Signaling in Mammalian CNS Angiogenesis and Blood-Brain Barrier Regulation. Neuron 95, 1056–1073 e1055 (2017).

26. Ulrich F, et al. Reck enables cerebrovascular development by promoting canonical Wnt signaling. Development 143, 147–159 (2016).

27. Vallon M, et al. A RECK-WNT7 Receptor-Ligand Interaction Enables Isoform-Specific Regulation of Wnt Bioavailability. Cell Rep 25, 339–349 e339 (2018).

28. Li H, et al. RECK in Neural Precursor Cells Plays a Critical Role in Mouse Forebrain Angiogenesis. iScience 19, 559–571 (2019).

29. Le V, et al. Mechanisms Underlying Rare Inherited Pediatric Retinal Vascular Diseases: FEVR, Norrie Disease, Persistent Fetal Vascular Syndrome. Cells 12, (2023).

30. Biswas S, et al. Glutamatergic neuronal activity regulates angiogenesis and blood-retinal barrier maturation via Norrin/beta-catenin signaling. Neuron 112, 1978–1996 e1976 (2024).

31. Chang J, et al. Gpr124 is essential for blood-brain barrier integrity in central nervous system disease. Nat Med 23, 450–460 (2017).

32. Bassett EA, et al. Norrin/Frizzled4 signalling in the preneoplastic niche blocks medulloblastoma initiation. Elife 5, (2016).

33. Wykoff CS, M. AMARONE Shows Promising Outcomes for a Novel Treatment Pathway in DME and nAMD. Retinal Physician 21, (2024).

34. Chidiac R, et al. A Norrin/Wnt surrogate antibody stimulates endothelial cell barrier function and rescues retinopathy. EMBO Mol Med 13, e13977 (2021).

35. Zhang L, et al. A Frizzled4-LRP5 agonist promotes blood-retina barrier function by inducing a Norrin-like transcriptional response. iScience 26, 107415 (2023).

36. Post Y, et al. Design principles and therapeutic applications of novel synthetic WNT signaling agonists. iScience 27, 109938 (2024).

37. O’Brien S, Chidiac R, Angers S. Modulation of Wnt-beta-catenin signaling with antibodies: therapeutic opportunities and challenges. Trends Pharmacol Sci 44, 354–365 (2023).

38. Nguyen H, Lee SJ, Li Y. Selective Activation of the Wnt-Signaling Pathway as a Novel Therapy for the Treatment of Diabetic Retinopathy and Other Retinal Vascular Diseases. Pharmaceutics 14, (2022).

39. Ding J, et al. Therapeutic blood-brain barrier modulation and stroke treatment by a bioengineered FZD(4)-selective WNT surrogate in mice. Nat Commun 14, 2947 (2023).

40. Martin M, et al. Engineered Wnt ligands enable blood-brain barrier repair in neurological disorders. Science 375, eabm4459 (2022).

41. Paszty K, et al. Plasma membrane Ca(2)(+)-ATPases can shape the pattern of Ca(2)(+) transients induced by store-operated Ca(2)(+) entry. Sci Signal 8, ra19 (2015).

42. Okunade GW, et al. Targeted ablation of plasma membrane Ca2+-ATPase (PMCA) 1 and 4 indicates a major housekeeping function for PMCA1 and a critical role in hyperactivated sperm motility and male fertility for PMCA4. J Biol Chem 279, 33742–33750 (2004).

43. Krebs J. Structure, Function and Regulation of the Plasma Membrane Calcium Pump in Health and Disease. Int J Mol Sci 23, (2022).

44. Stafford N, Wilson C, Oceandy D, Neyses L, Cartwright EJ. The Plasma Membrane Calcium ATPases and Their Role as Major New Players in Human Disease. Physiol Rev 97, 1089–1125 (2017).

45. Beckmann D, et al. Ca(2+) Homeostasis by Plasma Membrane Ca(2+) ATPase (PMCA) 1 Is Essential for the Development of DP Thymocytes. Int J Mol Sci 24, (2023).

46. Korthals M, et al. Plasma membrane Ca(2+) ATPase 1 (PMCA1) but not PMCA4 is critical for B-cell development and Ca(2+) homeostasis in mice. Eur J Immunol 51, 594–602 (2021).

47. Korthals M, et al. A complex of Neuroplastin and Plasma Membrane Ca(2+) ATPase controls T cell activation. Sci Rep 7, 8358 (2017).

48. Dalal PJ, Muller WA, Sullivan DP. Endothelial Cell Calcium Signaling during Barrier Function and Inflammation. Am J Pathol 190, 535–542 (2020).

49. Savage AM, et al. tmem33 is essential for VEGF-mediated endothelial calcium oscillations and angiogenesis. Nat Commun 10, 732 (2019).

50. Isshiki M, et al. Endothelial Ca2+ waves preferentially originate at specific loci in caveolin-rich cell edges. Proc Natl Acad Sci U S A 95, 5009–5014 (1998).

51. Hong SG, et al. Mechanosensitive membrane domains regulate calcium entry in arterial endothelial cells to protect against inflammation. J Clin Invest 134, (2024).

52. Dragoni S, Moccia F, Bootman MD. The Roles of Transient Receptor Potential (TRP) Channels Underlying Aberrant Calcium Signaling in Blood-Retinal Barrier Dysfunction. Cold Spring Harb Perspect Biol 17, (2025).

53. Moccia F, Negri S, Shekha M, Faris P, Guerra G. Endothelial Ca(2+) Signaling, Angiogenesis and Vasculogenesis: just What It Takes to Make a Blood Vessel. Int J Mol Sci 20, (2019).

54. Njegic A, Swiderska A, Marris C, Armesilla AL, Cartwright EJ. Plasma membrane calcium ATPase 1 regulates human umbilical vein endothelial cell angiogenesis and viability. J Mol Cell Cardiol 156, 79–81 (2021).

55. Long Y, et al. ATP2B1 Gene Silencing Increases NO Production Under Basal Conditions Through the Ca(2+)/calmodulin/eNOS Signaling Pathway in Endothelial Cells. Hypertens Res 41, 246–252 (2018).

56. Baggott RR, et al. Plasma membrane calcium ATPase isoform 4 inhibits vascular endothelial growth factor-mediated angiogenesis through interaction with calcineurin. Arterioscler Thromb Vasc Biol 34, 2310–2320 (2014).

57. Roux KJ, Kim DI, Raida M, Burke B. A promiscuous biotin ligase fusion protein identifies proximal and interacting proteins in mammalian cells. J Cell Biol 196, 801–810 (2012).

58. Zhang C, Lai MB, Khandan L, Lee LA, Chen Z, Junge HJ. Norrin-induced Frizzled4 endocytosis and endo-lysosomal trafficking control retinal angiogenesis and barrier function. Nat Commun 8, 16050 (2017).

59. Junge HJ. Ligand-Selective Wnt Receptor Complexes in CNS Blood Vessels: RECK and GPR124 Plugged In. Neuron 95, 983–985 (2017).

60. Cho C, Wang Y, Smallwood PM, Williams J, Nathans J. Dlg1 activates beta-catenin signaling to regulate retinal angiogenesis and the blood-retina and blood-brain barriers. Elife 8, (2019).

61. Hering H, Sheng M. Direct interaction of Frizzled-1, -2, -4, and -7 with PDZ domains of PSD-95. FEBS Lett 521, 185–189 (2002).

62. D’Angelo G, et al. GRASP65 and GRASP55 sequentially promote the transport of C-terminal valine-bearing cargos to and through the Golgi complex. J Biol Chem 284, 34849–34860 (2009).

63. Tauriello DV, et al. Wnt/beta-catenin signaling requires interaction of the Dishevelled DEP domain and C terminus with a discontinuous motif in Frizzled. Proc Natl Acad Sci U S A 109, E812–820 (2012).

64. Jho EH, Zhang T, Domon C, Joo CK, Freund JN, Costantini F. Wnt/beta-catenin/Tcf signaling induces the transcription of Axin2, a negative regulator of the signaling pathway. Mol Cell Biol 22, 1172–1183 (2002).

65. Hartman GD, et al. Ref-1 redox activity modulates canonical Wnt signaling in endothelial cells. Redox Biol 83, 103646 (2025).

66. Stamos JL, Weis WI. The beta-catenin destruction complex. Cold Spring Harb Perspect Biol 5, a007898 (2013).

67. Stahl A, et al. The mouse retina as an angiogenesis model. Invest Ophthalmol Vis Sci 51, 2813–2826 (2010).

68. Nitta T, et al. Size-selective loosening of the blood-brain barrier in claudin-5-deficient mice. J Cell Biol 161, 653–660 (2003).

69. Crabtree GR, Olson EN. NFAT signaling: choreographing the social lives of cells. Cell 109 Suppl, S67-79 (2002).

70. Suehiro J, et al. Genome-wide approaches reveal functional vascular endothelial growth factor (VEGF)-inducible nuclear factor of activated T cells (NFAT) c1 binding to angiogenesis-related genes in the endothelium. J Biol Chem 289, 29044–29059 (2014).

71. Huang T, Xie Z, Wang J, Li M, Jing N, Li L. Nuclear factor of activated T cells (NFAT) proteins repress canonical Wnt signaling via its interaction with Dishevelled (Dvl) protein and participate in regulating neural progenitor cell proliferation and differentiation. J Biol Chem 286, 37399–37405 (2011).

72. Ye X, Wang Y, Nathans J. The Norrin/Frizzled4 signaling pathway in retinal vascular development and disease. Trends Mol Med 16, 417–425 (2010).

73. Negri S, Faris P, Moccia F. Reactive Oxygen Species and Endothelial Ca(2+) Signaling: Brothers in Arms or Partners in Crime? Int J Mol Sci 22, (2021).

74. Shlobin NA, Har-Even M, Itsekson-Hayosh Z, Harnof S, Pick CG. Role of Thrombin in Central Nervous System Injury and Disease. Biomolecules 11, (2021).

75. Marceau F, et al. Bradykinin receptors: Agonists, antagonists, expression, signaling, and adaptation to sustained stimulation. Int Immunopharmacol 82, 106305 (2020).

76. Mugisho OO, Robilliard LD, Nicholson LFB, Graham ES, O’Carroll SJ. Bradykinin receptor-1 activation induces inflammation and increases the permeability of human brain microvascular endothelial cells. Cell Biol Int 44, 343–351 (2020).

77. Saneyoshi T, Kume S, Amasaki Y, Mikoshiba K. The Wnt/calcium pathway activates NF-AT and promotes ventral cell fate in Xenopus embryos. Nature 417, 295–299 (2002).

78. Park H, et al. Integrin-linked kinase controls retinal angiogenesis and is linked to Wnt signaling and exudative vitreoretinopathy. Nat Commun 10, 5243 (2019).

79. Vinyoles M, et al. Multivesicular GSK3 sequestration upon Wnt signaling is controlled by p120-catenin/cadherin interaction with LRP5/6. Mol Cell 53, 444–457 (2014).

80. Taelman VF, et al. Wnt signaling requires sequestration of glycogen synthase kinase 3 inside multivesicular endosomes. Cell 143, 1136–1148 (2010).

81. Yang M, et al. CTNND1 variants cause familial exudative vitreoretinopathy through the Wnt/cadherin axis. JCI Insight 7, (2022).

82. Boye K, et al. Endothelial Unc5B controls blood-brain barrier integrity. Nat Commun 13, 1169 (2022).

83. Wang Y, et al. Ephrin-B2 controls VEGF-induced angiogenesis and lymphangiogenesis. Nature 465, 483–486 (2010).

84. Roux KJ, Kim DI, Burke B, May DG. BioID: A Screen for Protein-Protein Interactions. Curr Protoc Protein Sci 91, 19 23 11–19 23 15 (2018).

