## Supplementary material for "The Calcium Pump ATP2B1/PMCA1 Regulates CNS Vascular Development by Facilitating Norrin- and WNT7A/B-induced Frizzled4 Signaling": Jo et al. 2026 supplement

**Supplementary Figures**

**A**

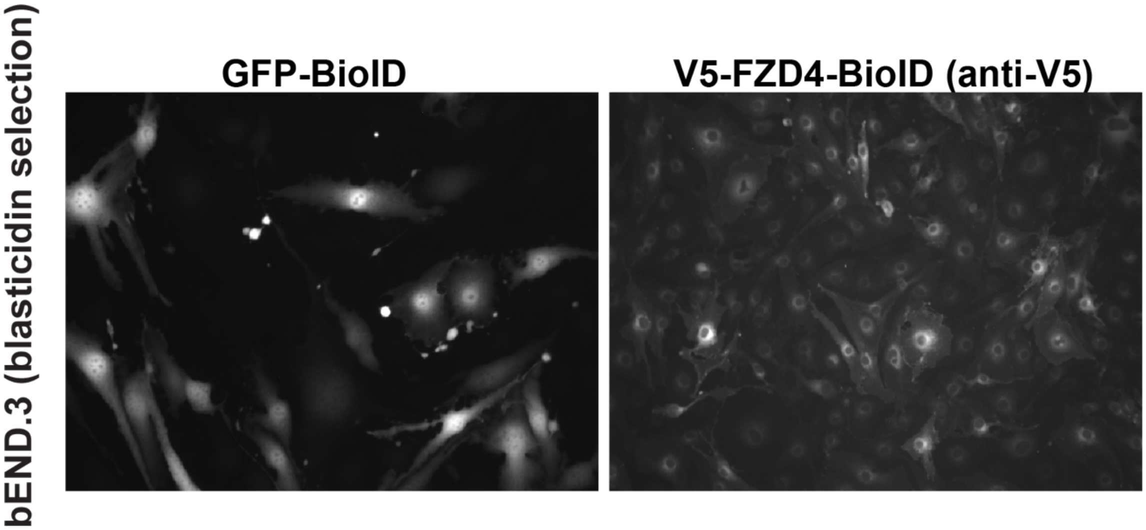

**Supplementary Fig. S1. Localization of GFP-BioID and V5-FZD4-BioID in stable bEnd.3 populations.** (**A**) GFP fluorescence was directly imaged, V5-FZD4-BioID was imaged after staining with anti-V5-Alexa488. V5-FZD4-BioID showed a perinuclear and plasma membrane localization, whereas GFP-BioID was broadly localized within the cells.

**A**

| Gene names | Sequence coverage [%], V5-FZD4-BioID | Abundance ratio (FZD4/GFP) | MS/MS count ratio (FZD4/GFP) | MS/MS count V5-FZD4-BioID (sum) |
|---|---|---|---|---|
| DLG1 | 50.3 | #DIV/0! | #DIV/0! | 105 |
| TSPAN12 | 29.8 | 175.88 | 13.67 | 84 |
| CTNND1 | 35.2 | #DIV/0! | 8.75 | 61 |
| FZD4 | 28.3 | #DIV/0! | #DIV/0! | 65 |
| SNAP23 | 81 | 50.49 | 14.50 | 48 |
| UBXN4 | 31.9 | 11.14 | 9.67 | 58 |
| SLC12A2 | 20.6 | #DIV/0! | #DIV/0! | 41 |
| VANGL1 | 35.9 | #DIV/0! | #DIV/0! | 56 |
| ADD3 | 28.3 | #DIV/0! | #DIV/0! | 41 |
| MCM2 | 33.2 | #DIV/0! | 26.00 | 47 |
| IMMT | 29 | #DIV/0! | #DIV/0! | 42 |
| ACBD3 | 40 | 12.56 | 8.33 | 45 |
| GORASP2 | 34.3 | #DIV/0! | 22.00 | 40 |
| ITGB1 | 20.6 | #DIV/0! | #DIV/0! | 33 |
| OCLN | 18 | #DIV/0! | #DIV/0! | 34 |
| PGRMC2 | 43.9 | 40.47 | 5.67 | 43 |
| DSC2 | 28 | #DIV/0! | 16.00 | 32 |
| STEAP3 | 23.2 | #DIV/0! | #DIV/0! | 21 |
| PTPN1 | 34.9 | #DIV/0! | #DIV/0! | 36 |
| DVL1 | 33.1 | #DIV/0! | 5.00 | 27 |
| VAPA | 48.2 | #DIV/0! | 15.00 | 27 |
| ATP2B1 | 15 | #DIV/0! | #DIV/0! | 28 |
| MARCKS | 36.7 | #DIV/0! | #DIV/0! | 21 |
| VANGL2 | 22.5 | #DIV/0! | #DIV/0! | 31 |
| CXADR | 40.3 | #DIV/0! | #DIV/0! | 23 |
| CDKAL1 | 28.3 | #DIV/0! | #DIV/0! | 30 |
| FNDC3A | 18 | #DIV/0! | #DIV/0! | 25 |
| LMAN2 | 26.1 | #DIV/0! | 6.50 | 23 |
| PVRL3 | 20 | #DIV/0! | #DIV/0! | 15 |
| STAM | 17 | #DIV/0! | 13.00 | 22 |
| CISD2 | 33.3 | #DIV/0! | 6.00 | 23 |
| DHRS7 | 36.3 | #DIV/0! | 6.00 | 26 |
| EFNB1 | 27.7 | #DIV/0! | 11.00 | 17 |
| NUMB | 23 | #DIV/0! | 11.00 | 17 |
| STX5 | 24.5 | #DIV/0! | 5.50 | 25 |
| RALA | 37.9 | #DIV/0! | 5.50 | 17 |
| TOMM40 | 33.5 | #DIV/0! | 5.50 | 23 |
| PVRL2 | 17.5 | #DIV/0! | #DIV/0! | 18 |
| EIF3H | 26.7 | #DIV/0! | 5.50 | 18 |
| LAMTOR1 | 74.5 | #DIV/0! | 11.00 | 20 |
| RABL3 | 45.3 | #DIV/0! | #DIV/0! | 13 |
| ATP6AP2 | 22.9 | #DIV/0! | #DIV/0! | 21 |
| MARCKSL1 | 48.2 | #DIV/0! | #DIV/0! | 14 |
| PTGES2 | 27.6 | #DIV/0! | 8.00 | 20 |
| UBIAD1 | 22.5 | #DIV/0! | #DIV/0! | 11 |
| NT5DC2 | 20.8 | #DIV/0! | 2.33 | 19 |
| CD99 | 15.7 | #DIV/0! | 6.00 | 16 |
| STX7 | 18 | #DIV/0! | 5.00 | 10 |
| BYSL | 15.3 | #DIV/0! | 4.00 | 14 |
| TBL2 | 15.2 | #DIV/0! | #DIV/0! | 11 |

**Supplementary Fig. S2. Proteins from 293T cells after bioinformatic filtering.** Note that when a value in GFP-BioID samples was 0, the ratio of FZD4/GFP could not be calculated, this is indicated with the error message #DIV/0!. Proteins of interest are highlighted in yellow markup.

**A**

| Gene names | Sequence coverage [%], V5-FZD4-BioID | Abundance ratio (FZD4/GFP) | MS/MS count ratio (FZD4/GFP) | MS/MS count V5-FZD4-BioID |
|---|---|---|---|---|
| CTNND1 | 54.5 | #DIV/0! | #DIV/0! | 85 |
| MYOF | 35.1 | 10.76 | 6.29 | 88 |
| ITGB1 | 38.5 | 23.87 | 6.67 | 85 |
| KTN1 | 40.7 | 14.23 | 5.43 | 86 |
| FZD4 | 21.4 | #DIV/0! | #DIV/0! | 76 |
| EGFR | 34.1 | #DIV/0! | #DIV/0! | 53 |
| ERBB2IP | 29.7 | 21.35 | 9.00 | 49 |
| POR | 52.1 | #DIV/0! | #DIV/0! | 56 |
| ATP2B1 | 21 | #DIV/0! | #DIV/0! | 43 |
| TOR1AIP1 | 47.5 | #DIV/0! | 22.00 | 49 |
| SLC1A5 | 31.6 | 24.31 | 5.50 | 47 |
| PPFIBP1 | 34.5 | #DIV/0! | #DIV/0! | 36 |
| FNDC3A | 28 | #DIV/0! | #DIV/0! | 41 |
| EPB41L1 | 33.8 | #DIV/0! | 21.00 | 35 |
| CDKAL1 | 58.9 | #DIV/0! | #DIV/0! | 36 |
| VANGL1 | 36.1 | #DIV/0! | 19.00 | 41 |
| PDXDC1 | 45.1 | #DIV/0! | #DIV/0! | 34 |
| ABCC1 | 21.6 | #DIV/0! | #DIV/0! | 35 |
| NUP155 | 21.2 | 10.43 | 6.00 | 40 |
| CXADR | 43.3 | #DIV/0! | #DIV/0! | 37 |
| SLC30A1 | 35.9 | 35.78 | 8.50 | 37 |
| ACBD3 | 45.1 | #DIV/0! | #DIV/0! | 31 |
| YKT6 | 66.7 | #DIV/0! | 16.00 | 34 |
| PGRMC2 | 43.9 | 19.97 | 5.33 | 41 |
| ADD3 | 36.7 | #DIV/0! | 15.00 | 31 |
| NSDHL | 51.7 | 27.81 | 7.50 | 37 |
| PTPN1 | 47.4 | 28.30 | 7.50 | 38 |
| ATP6AP2 | 34.6 | #DIV/0! | #DIV/0! | 24 |
| SNAP23 | 64.9 | #DIV/0! | #DIV/0! | 28 |
| LBR | 20.8 | 25.72 | 6.50 | 26 |
| CYB5A | 49.3 | #DIV/0! | #DIV/0! | 26 |
| GORASP2 | 30.8 | #DIV/0! | 12.00 | 27 |
| UBXN4 | 28.7 | #DIV/0! | 12.00 | 24 |
| STBD1 | 61.2 | #DIV/0! | #DIV/0! | 21 |
| GNAI1 | 53.4 | #DIV/0! | #DIV/0! | 17 |
| HMOX2 | 40.5 | #DIV/0! | #DIV/0! | 23 |
| VASN | 20.7 | #DIV/0! | #DIV/0! | 18 |
| SLITRK4 | 20.7 | #DIV/0! | #DIV/0! | 24 |
| VAPB | 45.7 | 19.09 | 5.00 | 26 |
| MARCKS | 43.4 | #DIV/0! | #DIV/0! | 13 |
| DLG1 | 26.8 | #DIV/0! | 9.00 | 17 |
| BET1 | 56.8 | #DIV/0! | 6.00 | 16 |
| CYB5B | 31.5 | #DIV/0! | #DIV/0! | 11 |

**Supplementary Fig. S3. Proteins from HeLa cells after bioinformatic filtering.** Note that when a value in GFP-BioID samples was 0, the ratio of FZD4/GFP could not be calculated, this is indicated with the error message #DIV/0!. Proteins of interest are highlighted in yellow markup.

**A**

| Gene names | Sequence coverage [%], V5-FZD4-BioID | Abundance ratio (FZD4/GFP) | peptide precursor abundance ratio (FZD4/GFP) | peptide precursor abundance (sum) |
|---|---|---|---|---|
| Mcam | 46.00 | 17.49 | 4.00 | 60.00 |
| Ano6 | 25.00 | 10.79 | 5.71 | 40.00 |
| Hmox2 | 65.00 | 14.22 | #DIV/0! | 39.00 |
| Thsd1 | 32.00 | 23.19 | 12.67 | 38.00 |
| Tfrc | 32.00 | 8.51 | 9.00 | 36.00 |
| Pecam1 | 37.00 | 7.72 | 6.00 | 36.00 |
| Tmpo | 46.00 | 5.79 | 2.92 | 35.00 |
| Itgb1 | 23.00 | 7.09 | 3.20 | 32.00 |
| Lbr | 27.00 | 5.43 | 3.10 | 31.00 |
| Ephb4 | 18.00 | 5.42 | 7.50 | 30.00 |
| Esam | 44.00 | 10.28 | 3.63 | 29.00 |
| Eif2ak3 | 19.00 | 5.39 | 29.00 | 29.00 |
| Tgfbr2 | 30.00 | 9.68 | #DIV/0! | 28.00 |
| Dlg1 | 29.00 | 7.52 | 7.00 | 28.00 |
| Sema4c | 20.00 | 14.47 | 26.00 | 26.00 |
| Fzd4 | 16.00 | 100.00 | #DIV/0! | 22.00 |
| Unc5b | 20.00 | 13.44 | #DIV/0! | 22.00 |
| Pgrmc2 | 55.00 | 51.51 | 7.00 | 21.00 |
| Atp2b1 | 16.00 | 6.56 | #DIV/0! | 20.00 |
| Pvrl2; Nectin2 | 28.00 | 35.75 | #DIV/0! | 19.00 |
| Vangl1 | 30.00 | 23.57 | #DIV/0! | 18.00 |
| Slc3a2 | 18.00 | 5.60 | 18.00 | 18.00 |
| Arhgef15 | 19.00 | 5.13 | 9.00 | 18.00 |
| Slc1a5 | 20.00 | 20.71 | #DIV/0! | 17.00 |
| Dvl1 | 19.00 | 5.75 | 8.50 | 17.00 |
| Cdh5 | 17.00 | 5.36 | 4.25 | 17.00 |
| Slc25a24 | 27.00 | 5.21 | #DIV/0! | 17.00 |
| Acbd3 | 24.00 | 6.19 | #DIV/0! | 16.00 |
| Snap23 | 52.00 | 9.54 | 6.50 | 13.00 |
| Tmem199 | 31.00 | 28.06 | 12.00 | 12.00 |
| Thbd | 19.00 | 21.60 | #DIV/0! | 12.00 |
| Gorasp2 | 14.00 | 7.99 | #DIV/0! | 11.00 |
| Cdc42ep1 | 19.00 | 23.76 | #DIV/0! | 10.00 |
| Jam2 | 21.00 | 5.47 | #DIV/0! | 10.00 |
| Vamp3 | 64.00 | 8.98 | 9.00 | 9.00 |
| Cyb5b | 53.00 | 8.18 | #DIV/0! | 9.00 |
| Palm | 14.00 | 6.57 | #DIV/0! | 9.00 |
| Tmem51 | 23.00 | 17.97 | #DIV/0! | 8.00 |
| Ckm | 17.00 | 100.00 | #DIV/0! | 7.00 |
| Cd200 | 16.00 | 13.38 | #DIV/0! | 6.00 |
| Spry4 | 18.00 | 8.21 | #DIV/0! | 6.00 |
| Vamp5 | 31.00 | 26.71 | #DIV/0! | 5.00 |
| Cisd2 | 27.00 | 6.40 | #DIV/0! | 5.00 |
| Hprt | 17.00 | 100.00 | #DIV/0! | 4.00 |
| Myl1 | 30.00 | 100.00 | #DIV/0! | 3.00 |
| Marcksl1 | 14.00 | 6.95 | 3.00 | 3.00 |

**Supplementary Fig. S4. Proteins from bEnd.3 cells after bioinformatic filtering.** Note that when a value in GFP-BioID samples was 0, the ratio of FZD4/GFP could not be calculated, this is indicated with the error message #DIV/0!. Proteins of interest are highlighted in yellow markup.

A

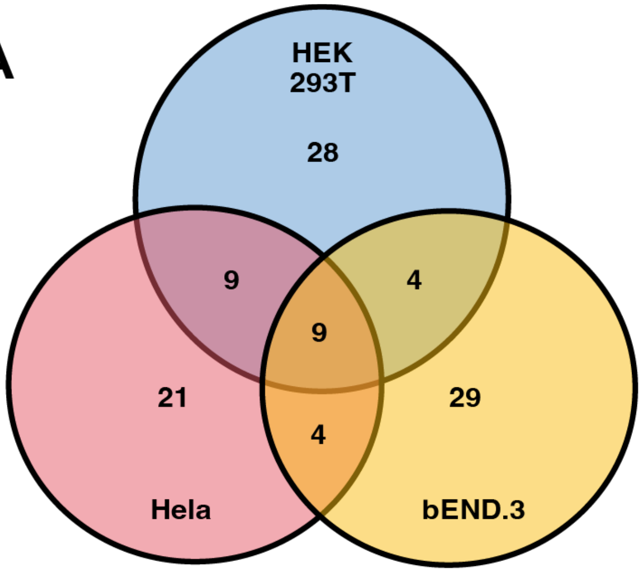

B

ABCD3
ATP2B1
CTNND1
DLG1
FZD4
GORASP2
ITGB1
PGRMC2
SNAP23
VANGL1

**Supplementary Fig. S5. Overlap in the identified proteins from three cell lines after bioinformatic filtering.** (**A**) Venn diagram indicates partial overlap of filtered proteins between two or three cell lines. (**B**) List of proteins identified in all three cell lines in alphabetical order.

A

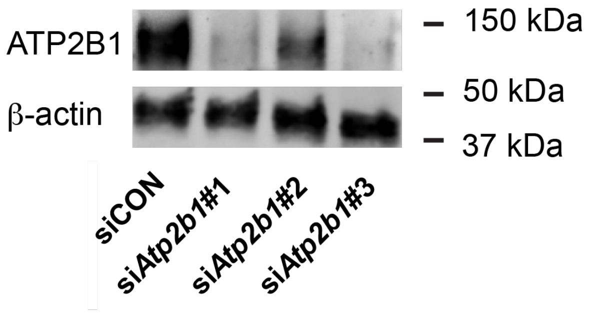

B

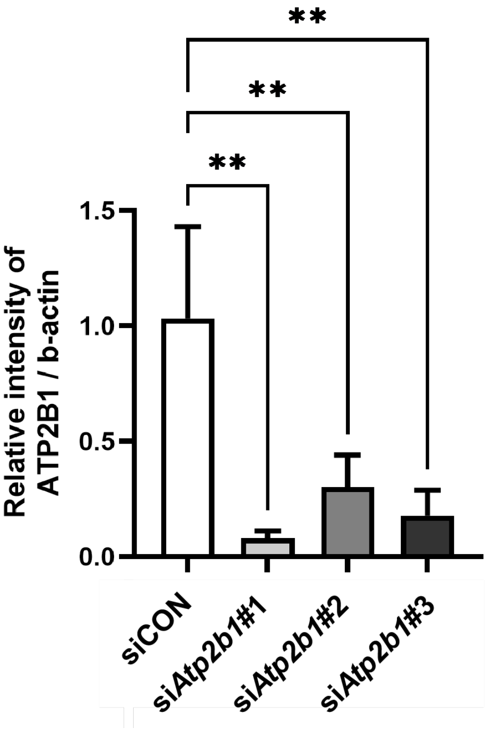

**Supplementary Fig. S6. ATP2B1 protein knock down efficiency.** (**A**) siRNAs at 20 nM final concentration knocked down mATP2B1 protein 48 hours post-transfection in bEnd.3 cells. (**B**) Quantification of band intensity, n=4-6 replicates, mean +/- SD shown, one-way ANOVA with Dunett's post hoc.

A

| Name | minus Norrin RPKM | plus Norrin RPKM | Fold change | heteroscedastic TTEST |
|---|---|---|---|---|
| Axin2 | 0.123 | 2.63 | 21.3178682 | 0.000554 |
| P2ry1 | 2.359 | 13.039 | 5.52667651 | 0.000009 |
| Atp2b1 | 40.656 | 35.174 | 0.86517966 | 0.174234 |
| Atp2b2 | 0.515 | 0.403 | 0.78170182 | 0.213296 |
| Atp2b3 not detected | | | | |
| Atp2b4 | 0.472 | 0.404 | 0.8563397 | 0.898660 |
| Nfatc1 | 23.943 | 19.387 | 0.80972806 | 0.039231 |
| Nfatc2 | 0.351 | 1.435 | 4.08440847 | 0.006796 |
| Nfatc3 | 22.923 | 21.623 | 0.94331346 | 0.674988 |
| Nfatc4 | 0.428 | 1.189 | 2.78141267 | 0.010636 |
| Nfat5 | 95.947 | 120.258 | 1.25338593 | 0.029474 |
| Fzd4 | 51.303 | 46.953 | 0.91520812 | 0.359114 |
| Lrp5 | 102.579 | 85.754 | 0.83597757 | 0.060010 |
| Lrp6 | 29.999 | 31.056 | 1.03524102 | 0.404175 |
| Adgra2 (GPR124) | 97.907 | 95.934 | 0.9798465 | 0.993949 |
| Reck | 2.803 | 2.645 | 0.94373397 | 0.612772 |

**Supplementary Fig. S7. Selected transcripts identified from bulk-RNAseq in bEnd.3 cells.** Quadruplicate samples of bEnd.3 cells were stimulated with vehicle or Norrin and subjected to bulk RNA-seq. *P2ry1* was identified as a response gene. The table shows that *Atp2b1* is the predominant paralog in bEnd.3 cells.

**A**

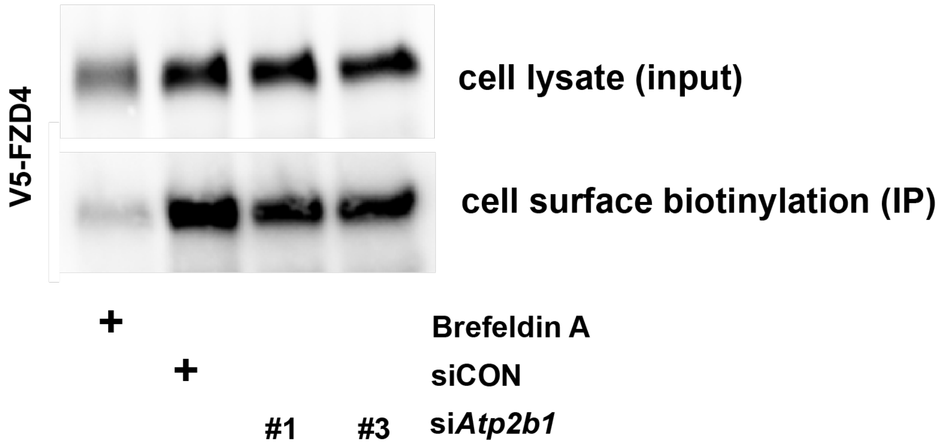

**B**

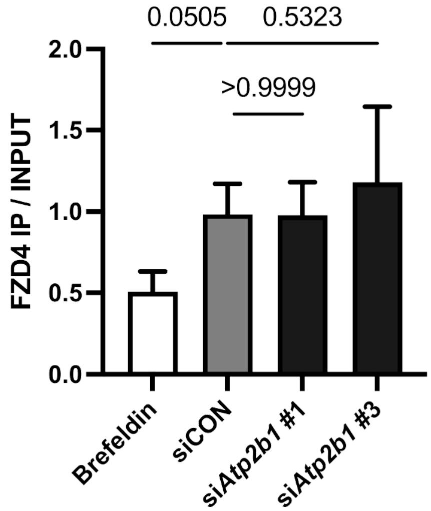

**Supplementary Fig. S8. *Atp2b1* siRNA does not significantly alter FZD4 cell surface steady state.** (**A**) A stably transduced bEnd.3 cell population expressing V5-FZD4 was subjected to cell-surface biotinylation. The amount of FZD4 in the lysate and after Neutravidin pull down was determined with antibody directed against the V5 tag. (**B**) Quantification of band intensity from three independent experiments. Mean +/- SD shown, one-way ANOVA with Tukey's post hoc.

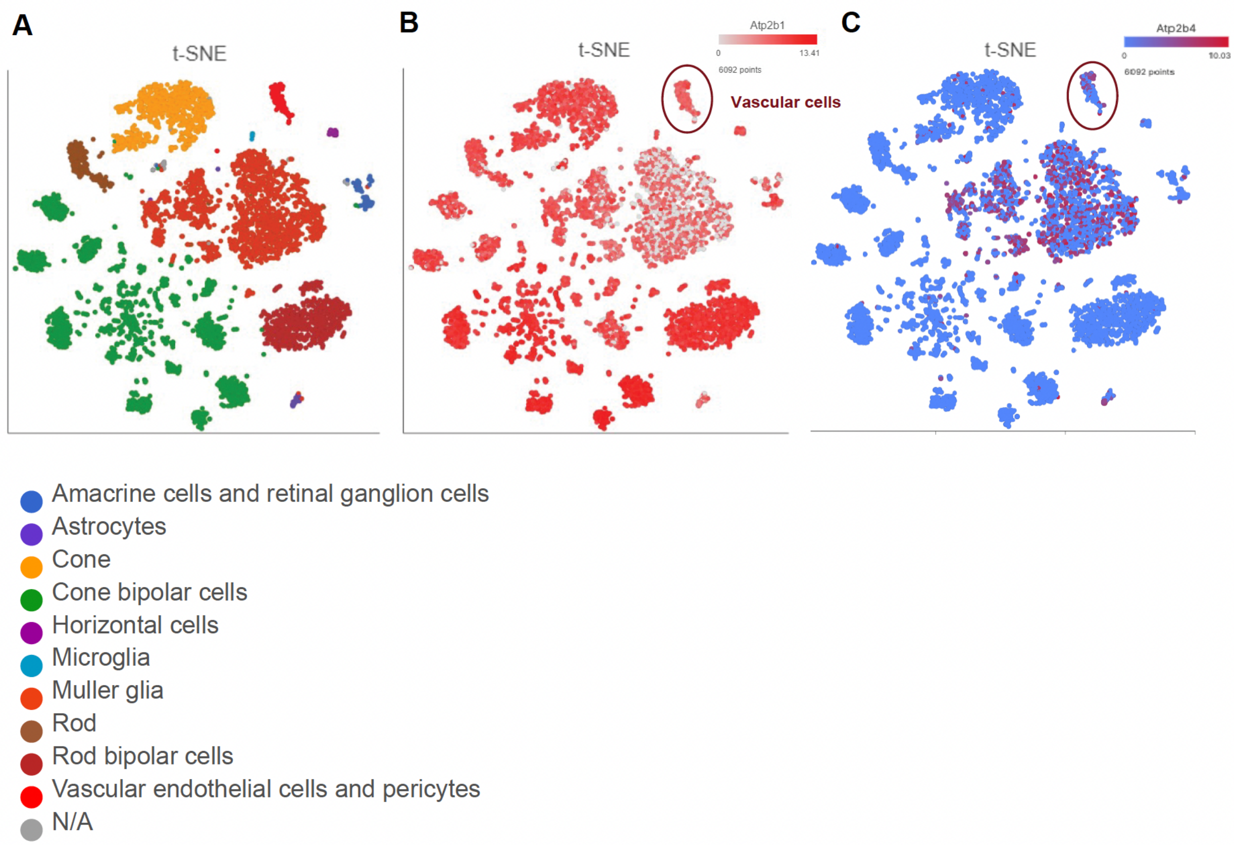

**Supplementary Fig. S9. Re-analysis of published scRNAseq data.** Our data in Zhang et al. 2023 show broad expression of *Atp2b1* in multiple retinal cell types, including vascular cells, and more restricted expression of *Atp2b4*, including in vascular cells (vascular cells highlighted by oval frame).

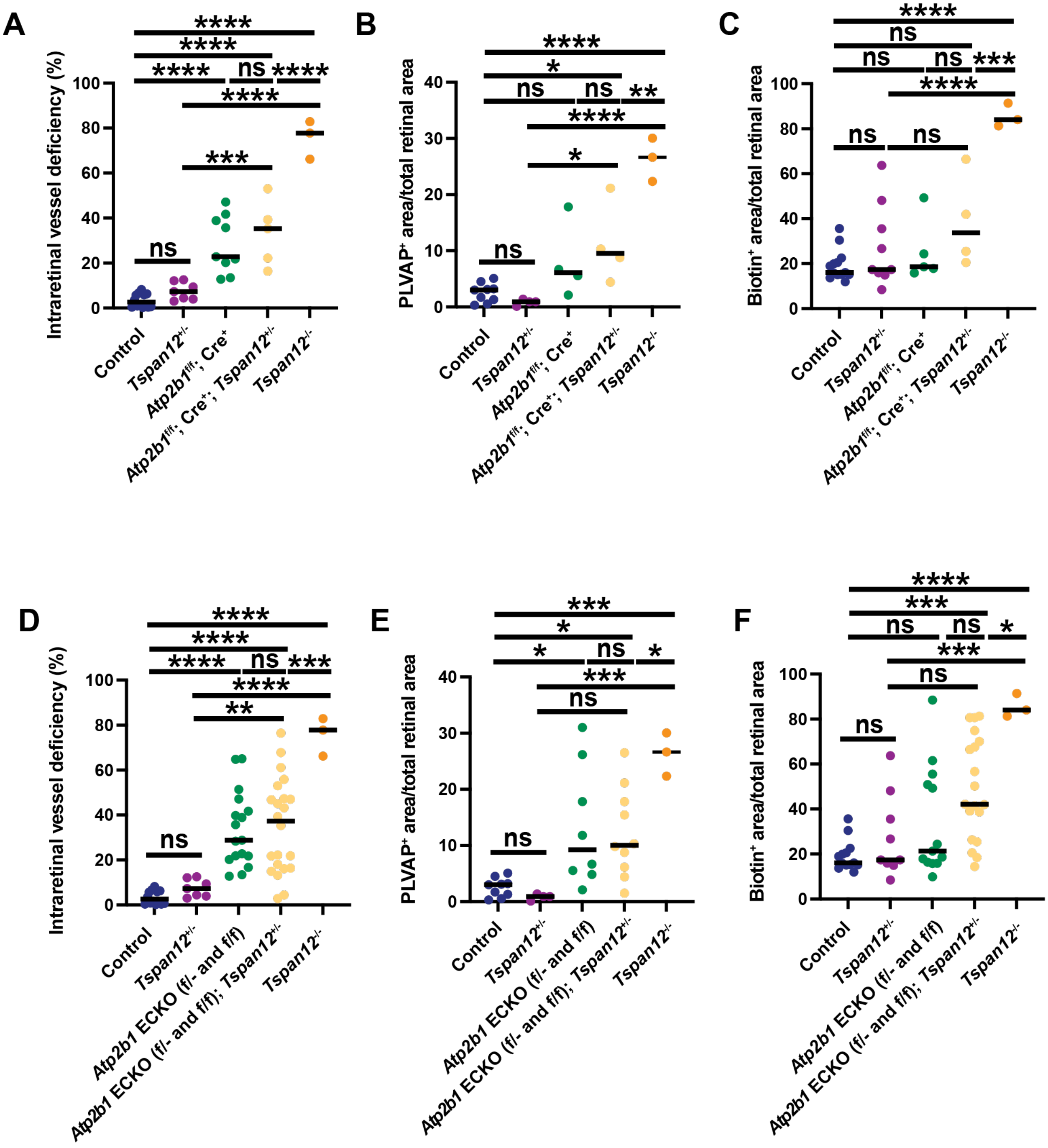

**Supplementary Fig. S10. Supplementary analysis of BRB phenotypes in *Atp2b1* ECKO mice.** (**A-C**) Disaggregated analysis of *Atp2b1*$^{f/f}$; Cre$^{+}$ ECKO allele series. n=3-14 retinas from 3-14 mice, graphs represent median, one-way ANOVA with Tukey's post hoc. (**D-E**) Aggreagted analysis of the *Atp2b1*$^{f/f}$; Cre$^{+}$ and *Atp2b1*$^{f/f}$; Cre$^{+}$ ECKO allele series combined. n=3-22 retinas from 3-22 mice, graphs represent median, one-way ANOVA with Tukey's post hoc.

**Supplementary methods**

ORBITRAP FUSION LIQUID CHROMATOGRAPHY-MASS SPECTROMETRY.

To analyze peptides from bEnd.3 cells, we reconstituted the dried peptide mixtures in 98:2:0.1,H2O:acetonitrile (ACN):formic acid (FA) and analyzed ~200 nanogram of each sample by capillary LC-MS on an Orbitrap Fusion mass spectrometer (Thermo Fisher Scientific, Inc., Waltham, MA) online with a Thermo UltiMate™ 3000 RSLCnano LC system. Peptides were separated on a 40 cm self-packed C18 capillary column with 100 um inner diameter, with Dr. Maisch GmbH ReproSil-PUR 12 Å C18-AQ, 1.9 um particle size; the column was maintained at 55°C with a column heater from Sonation (Biberach, Germany). Peptides were loaded directly on column at 325 nl/min with 98:2:0.1,H2O:ACN:FA. We performed elution of the peptides with the following gradient: 5 – 32% solvent B from 0 – 50 minutes, 32 – 45% solvent B from 50 – 55 minutes, 45 – 90% solvent B from 55 – 60 minutes 325 nl/minute, where solvent A was 0.1% formic acid in water and solvent B was 0.1% formic acid in ACN. We operated the mass spectrometer with the following parameters: ESI voltage 2.1kV, ion transfer tube 275 °C; Orbitrap MS1 scan 120k resolution in profile from 380 – 1580 m/z with 50 msec injection time and 120% normalized automatic gain control (AGC); MS2 triggered on precursors charge states 2 – 5 above 20000 counts; precursor isolation window 1.6 Da; MIPS (monoisotopic peak determination) set to Peptide; higher energy collisional dissociation (HCD) at 35% with MS2 Orbitrap detection at 50k resolution (at 200 m/z), 86 msec injection time,  100% (1000) AGC, dynamic exclusion duration 15 sec with +/- 10 ppm mass tolerance.

DATABASE SEARCHING.

We processed peptide tandem MS using SEQUEST (Thermo Scientific) in Proteome Discoverer 4.0. The mouse Universal Proteome database (UP000000589) was downloaded from UniProt on 11/19/2021 and merged with a common lab contaminant protein database (https://www.thegpm.org/crap/) (55,214 total protein sequences). We applied the precursor mass recalibration node with precursor mass tolerance 20 ppm, product ion tolerance 0.2 Da with fixed carbamidomethyl (CAM) modification of cysteine

57.0215 m/z. The SEQUEST (PMID: 24226387) database search parameters included enzyme trypsin full specificity, 2 missed cleave sites; precursor tolerance 15 ppm, fragment ion tolerance 0.05 Da and maximum 4 dynamic modifications per peptide. We specified CAM cysteine (+57.021 Da) as a fixed modification and the dynamic modifications were acetylation of protein N-terminus (+42.011 Da), oxidation of methionine (+15.995 Da), conversion of glutamine to pyroglutamic acid (-17.027 Da), methionine loss at the protein N-terminus (-131.040 Da), methionine loss + acetylation at the protein N-terminus (-89.030 Da), biotinylation of lysine (226.078 Da) and deamidation of asparagine or glutamine (+0.984 Da).

CRITERIA FOR PROTEIN IDENTIFICATION.

We applied 1% protein and peptide False Discovery Rate (FDR) filters using the Percolator algorithm (https://doi.org/10.1038/nmeth1113) in PD.

LABEL FREE QUANT

We used the label free quantification workflow in PD 2.4 that included steps for feature extraction, chromatographic alignment, peptide mapping to features, protein abundance calculation, normalization, protein relative abundance ratio calculation and hypothesis testing for significance of relative fold chance. We applied the Minora Feature Detector algorithm in PD 2.3 for the assignment of chromatographic features for isotopically related peaks within a 0.2 minute retention time range. After features were assigned, chromatographic alignment was performed; the file with the largest number of features was assigned as the reference to which features from each file were aligned within a 4 minute (RT) window tolerance and 12 ppm mass tolerance. Retention times were adjusted from a non-linear regression-based fit of the differences in retention time between reference and sample. Peptide were mapped to retention time-aligned consensus features across samples with the requirement that at least one sample contains a peptide spectral match. We applied total peptide summed normalization across samples. Protein abundance calculations were made from summed peptide abundances. We used the background t-test for hypothesis testing with the null hypothesis of equal protein abundance. Protein relative abundances between groups were reported with p-values adjusted with the Benjamini-Hochberg method for multiple testing corrections.

**Table S1. qPCR primers**

| qPCR primers | Sequence (5'-3') |
|---|---|
| m*Atp2b1* fw | TGTGTGTGGTGTTGGTGACGGC |
| m*Atp2b11* rev | TGCCCACCCCTGATGACGGT |
| m*Axin2* fw | CAGGAGGATGCTGAAGGCTC |
| m*Axin2* rev | GCAGGCAAATTCGTCACTCG |
| m*P2ry1* fw | CATGAAGCCTTGGAGCGGCA |
| m*P2ry1* rev | GAGGGCTGGTAGGGTGAGCA |
| m*Egr3* fw | CGCGCTCAACCTCTTCTCCG |
| m*Egr3* rev | TCACGGTCTTGTTGCCGGGG |
| m*Nr4a2* fw | CGGCCTGTCAGCACTACGGT |
| m*Nr4a2* rev | ACTGACAACGATTTCGGCGGC |

*m; mouse, fw; forward, rev; reverse*
